# Benchmarking normalisation methods for differential binding analysis in CUT&RUN

**DOI:** 10.64898/2026.05.24.727463

**Authors:** Krutika Ambani, Antonio Ahn, April C. Watt, Catherine Blyth, Brendan Russ, Michael Taylor, Shom Goel, Alicia Oshlack

## Abstract

CUT&RUN (*Cleavage Under Targets and Release Using Nuclease*) is an increasingly popular method for profiling protein interactions (transcription factors, histone modifications, etc) with DNA across the whole genome. When performing differential binding analysis of CUT&RUN data to identify genomic regions where interaction profiles vary between conditions, data normalisation is essential for accurate biological interpretations. Despite this, there are no clear guidelines on the optimal normalisation method for CUT&RUN datasets. Here, we examine five normalisation approaches (spike-in, library size, background, reads-in-peak and greenlist) and highlight that different methods can result in widely discrepant interpretations of the data. We test these normalisation methods by simulating a variety of plausible differential binding scenarios as well as in-house generated datasets. We determined normalisation by data driven methods to be the most robust. Importantly, we find spike-in normalisation to have variable performance across experiments likely due to additional stochastic variations in wet-lab procedures and suggest that their use should be justified with orthogonal assays. Our findings inform the use of normalisation methods for CUT&RUN data and should thus facilitate reproducible and robust analysis.

## Introduction

Proteins that interact with chromatin regulate the initiation, continuation, suppression and termination of gene transcription whilst also influencing DNA conformation(1). In differential analysis, the goal is to identify genomic regions where a protein’s interaction with the chromatin is significantly increased or decreased across different conditions(2). Characterising these regions can give insights into the activity of transcriptional programs, the structure of the chromatin and resulting phenotypes.

Cleavage Under Targets, Release Using Nuclease (CUT&RUN) is a recently developed method of whole-genome profiling of interactions of proteins (e.g. transcription factors, transcriptional cofactors, and histone tails bearing post-translational modifications) with DNA(3). This technique has become increasingly popular as it is able to profile DNA-protein interactions with higher signal-to-noise ratio, fewer number of input cells, and lower sequencing depth compared to previously available methods such as ChIP-seq(3).

However, as in any profiling technique, in CUT&RUN there are multiple sources of technical variation. These can broadly be categorised into those relating to the wet-lab and experimental protocol, and those relating to sequencing procedures. Technical variations in the wet-lab include variation in cell number between samples, sample loss, hand-to-hand differences between users, differences in digestion time/temperature and amplification bias. Technical variation due to sequencing technology could include differences in sequencing depth between samples and composition bias where contaminants or regions in the genome preferentially utilise the finite number of available sequencing reads leading to an apparent depletion in other regions of the genome which are not biologically relevant.

To remove the variation in the data due to the technical factors, it is critical that a robust method of normalisation is implemented(4). The method of normalisation chosen depends on the underlying assumptions, perceived technical biases, and the analysis being conducted(5).

The CUT&RUN protocol established by Skene et al. recommends the use of spike-in chromatin as the normalisation strategy for accurate quantitative assessment of protein occupancy(3). In that study, normalisation to chromatin spike-in revealed that the number of cleavage events scaled proportionally with the input cell number. However, in the context of differential binding analysis in CUT&RUN, researchers are interested in the average change in protein binding at genomic regions - averaged across cells within a sample - between treatment or biological conditions. In this context, the differences in cell number between samples would be an example of technical variation that an ideal normalisation method would eliminate. Further to this, the overall robustness of the spike-in method has been challenged in other differential analysis contexts including in RNA-Seq(6) and ChIP-Seq(7), calling into question the usability of spike-in normalisation in the context of CUT&RUN differential analysis.

While there have been comparisons of normalisation methods for ChIP-seq data(7–9), there are currently no clear guidelines as to the optimal method of normalisation in differential binding analysis in CUT&RUN. In this study, we evaluated the performance of five normalisation methods: library size, reads-in-peak, background, greenlist(10) and spike-in(9) (further detail of each method in the ‘Materials and Methods’ section) in both simulation studies and in-house generated datasets. We found that data-driven normalisation approaches as implemented in DiffBind are the most robust methods, while spike-in normalisation are only appropriate in some settings.

## Materials and Methods

### Cell line maintenance

Human breast cancer cell line MCF7 was obtained from the American Type Culture Collection (ATCC) and was cultured in RPMI-1640 medium (Gibco) supplemented with 10% fetal bovine serum (GE HyClone), 1% GlutaMax (Gibco) and 1% HEPES (Gibco) and incubated at 37°C and 5% CO_2_ in a humidified incubator. Cells were tested negative for mycoplasma by PCR and cell line identity was confirmed by short tandem repeat analysis (Promega GenePrint 10 System). Drosophila S2 cells were cultured in S2 medium (Gibco) supplemented with 10% heat inactivated FBS and cultured at room temperature.

### Drug treatment

Estradiol was obtained from Sigma Aldrich and dissolved in ethanol. For hormone deprivation and stimulation experiments, MCF7 cells were cultured in phenol-free RPMI medium supplemented with 1% GlutaMax, 1% HEPES, and charcoal-stripped serum for two to three days. Cells were stimulated with 1 nM estradiol for one hour. Following treatment, cells from the same culture were divided into separate samples for CUT&RUN, representing technical replicates.

Abemaciclib methanesulfonate was obtained from MedChemExpress and reconstituted in dimethyl sulfoxide (DMSO). MCF7 cells were plated into separate flasks to establish biological replicate cultures and independently treated with 500 nM abemaciclib or DMSO vehicle control for two days.

#### Gene knockout

Custom single-guide RNA were obtained from Integrated DNA Technologies (IDT). sgRNA and Alt-R^TM^ S.p. Cas9 Nuclease V3 (IDT) were nucleofected into MCF7 cells using the 4D-Nucleofector X Unit (Promega). gRNA sequence against *ESR1*: 5’ - CTGACCGTAGACCTGCGCGT; non-targeting control: 5’-CATTTCTCAGTGCTATAGAG.

### CUT&RUN

CUT&RUN was performed using the Cell Signaling Technologies (CST) CUT&RUN Assay Kit. All buffers were prepared as per manufacturer’s protocol unless otherwise specified. For each CUT&RUN reaction, 500,000 cells were used. For spike-in with Drosophila S2 cells, 20,000 cells were added to each reaction prior to the first wash step. Spike-in with yeast DNA fragments was performed as per manufacturer’s instructions. Briefly, spike-in DNA provided by the CUT&RUN Assay Kit was diluted 1 in 100 in nuclease-free water. Each CUT&RUN reaction received 1 μL of diluted spike-in DNA when Mnase digestion was ended by the STOP buffer. The STOP Buffer was modified, with 0.1% SDS, Proteinase K, and 300 mM NaCl added to release all DNA material. Samples were incubated at 55°C for 1 hour and DNA was extracted using the DNA Purification Buffers and Spin Columns kit (CST). AMPure XP beads (Beckman Coulter) were used to select DNA fragments that were 0-700 bp long. Concentration and fragment sizes of the DNA were measured using the TapeStation D1000 HS kit (Agilent).

CUT&RUN libraries were constructed using the NEBNext Ultra II Kit (New England Biolands; NEB) and Multiplex Oligos for Illumina (96 Unique Dual Index Primer Pairs (NEB), following the “CST DNA Library Prep Kit for Illumina Protocol” that was specifically optimised for CUT&RUN, which included modifications to adaptor dilutions, AMPure XP bead ratios, number of PCR amplification cycles, and the PCR anneal and extension step, which was shortened to 13 seconds. Amplified DNA was cleaned up using AMPure XP beads and eluted in 10 mM Tris pH 8.0. Libraries were then pooled and sequenced on a NextSeq 2000 instrument, with a goal of 5-10 million 100 bp read-pairs per sample.

Primary antibodies used include anti-BRG1 (CST #49360), anti-ER (CST #8644), rabbit IgG (CST #66362), and Active Motif Spike-in Antibody (AM61686).

#### Western Blotting

To detect estrogen receptor expression, cells were lysed using 1X laemmli buffer with 5% beta-mercaptoethanol. Lysates were boiled at 95°C for 13 minutes. To detect nuclear BRG1 protein levels, nuclei were extracted and lysed using the Nuclear Complex Co-IP kit (Active Motif) as previously described(11). Western blotting was performed as previously described(12). Ponceau S was used to stain the membrane for total protein and histones for loading control. The primary antibodies used were anti-ER (Abcam; ab3575) and anti-BRG1 (CST #49360).

### Raw sequencing data Processing

Read quality of all raw and trimmed FASTQ files was assessed with FastQC(13) (v0.12.1) and summarised with MultiQC(14) (v1.24.1).

#### CUT&RUN

The FASTQ files of the CTCF CUT&RUN dataset GSE84474 were downloaded from Gene Expression Omnibus (GEO). Raw data were processed using the following set of tools.

The composition of the raw FASTQ files was screened using FastQ Screen(15) (v0.15.3). Adaptors and poor-quality reads were trimmed for the BRG1 and ER datasets using bbduk.sh from BBmap(16) (v38.90) with parameters: ktrim=r k=23 mink=11 hdist=1 tpe tbo qtrim=r trimq=5 minlen=20.

In order to calculate spike-in normalization factors based on the total number of reads aligned to the spike-in genome, a hybrid genome approach was used. Hybrid genomes were created by concatenating the raw FASTA files of species present in the sample and indexed using bowtie2(17) (v2.3.4.1) bowtie2-build. For the BRG1 dataset, a hybrid reference genome was constructed by combining the human GRCh38(18) analysis set (GCA_000001405.15_GRCh38_no_alt_plus_hs38d1_analysis_set; NCBI) with the *Saccharomyces cerevisiae* S288C R64 reference genome(19) (GCF_000146045.2; NCBI RefSeq), and samples were aligned to this hybrid assembly. For the CTCF dataset, a hybrid genome was constructed from human GRCh38 and *Drosophila melanogaster* dm6(20) (BDGP Release 6; UCSC Genome Browser). For the ER dataset, a combined genome was constructed from the concatenation of the human GRCh38, *Drosophila melanogaster* dm6(20) and the *Saccharomyces cerevisiae* S288C R64 reference genome(19).

Alignment to the relevant hybrid genome was conducted with bowtie2 (v2.3.4.1) using parameters --dovetail --local --very-sensitive --no-mixed --no-discordant --phred33 -I 10 -X 700 -p 14. Mitochondrial and spike-in species reads were removed from the BAM files using samtools(21) (v1.9).

Duplicates were marked but not removed with Picard(22) (v3.0.0). Reads in the BAM files overlapping with the blacklisted regions from ChIP(23) were removed using BEDTools (v2.27.1).

Peaks were called using MACS2(24) (v2.2.7.1) callpeaks function with the parameters -f BAMPE and -g hs. For the BRG1 and ER datasets, a q-value threshold of 0.05 was applied, and an IgG input sample was provided as a control via the -c parameter. For the CTCF dataset, no IgG input sample was available, and a q-value threshold of 0.01 was used.

### Differential binding analysis

The Bioconductor package DiffBind^2^ in R was used to generate a consensus peak set (peak width of 401bp) and identify differentially bound sites (terms “peak” and “binding site” used interchangeably). Default parameters were used throughout, except in dba.normalize(). An FDR cut-off of 0.05.

### Overview of normalisation methods tested

#### Library size normalisation

Library size normalisation, the default method in DiffBind, calculates scale factors as:

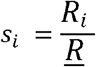

where *s_i_* is the scale factor for sample *i, R*_*i*_ is the total read count in the corresponding BAM file, and *R* is the mean total read count across all samples. This approach adjusts for differences in sequencing depth between samples.

#### Reads-in-peak normalisation

Reads-in-peak (RiP) normalisation computes scale factors using the median-of-ratios approach. A pseudo-reference is generated from the geometric mean of raw counts across all samples for each peak. Scale factors are calculated as:

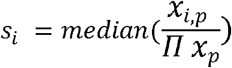

where *s_i_* is the scale factor for sample *i,x*_*i,p*_ is the raw counts for a peak *p* in sample *IIx_p_*, and is the geometric mean of the raw counts for peak *p* across all samples. By taking the median ratio, scaling is driven by the least variable peak across samples. This method was implemented using dba.normalize() with parameters normalize = RLE and library = RiP. RLE refers to “Relative Log Expression” described here (26).

#### Background normalisation

For background normalisation, chromosomes containing at least one consensus peak were divided into 15,000 bp bins. Reads from BAM files were tallied per bin to generate a counts matrix (bins × samples). Scale factors were derived assuming a core set of background bins remain unchanged across conditions:

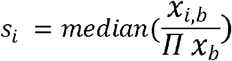

where *s_i_* is the scale factor for sample *i, x_i,b_* is the raw counts for a background bin *b* in sample *i*, and *Π x _b_* is the geometric mean of the raw counts for background bin *b* across all samples. Here, the scaling is driven by the least variable background bin across samples. Background normalisation was implemented with *normalize =* “*RLE*”, *library =* “*background*”, *background=TRUE* parameters within the dba.normalize() function.

#### Spike-in normalisation

The scale factors for spike-in normalisation were calculated as follows:

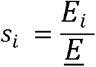

where *s_i_* is the scale factor for sample *i, E_i_* is the total number of reads in the spike-in BAM file for sample *i*, and *E* is the mean of the total number of spike-in reads of all samples in the experiment. Scale factors were manually specified per sample in the dba.normalize() function.

#### Greenlist normalisation

The scale factors calculated were based on the counts that overlap the regions of non-specific signal as per the get_sizeFactors.R script obtained from GitHub https://github.com/fndemello/CUT-RUN_greenlist(10) . These scale factors were specified by sample in the dba.normalize() function.

### Characterisation of peak distribution on chromatin

Genomic locations (i.e. gene promoters versus non-promoters) of transcription factor binding sites were determined using the annotatepeak() function in *ChIPseeker*^25^ *(v 1*.*42*.*0)*. Gene promoters were defined as the region -2000 to 500 base pairs from a transcription start site (TSS). Transcript annotations were obtained from the Gencode version 35 human basic annotations GTF file(27) .

Gene set enrichment testing of differentially bound peaks was conducted using the chipenrich() function from the ChIP-Enrich(28) (v 2.32.0) package. Genes were identified by the TSS nearest to each peak of interest. The Hallmark gene sets from the Human Molecular Signatures Database were used(29) .

### Generating datasets for Negative Experiment

The raw data of all 17 samples in the CTCF CUT&RUN dataset (GSE84474) were initially considered (Supplementary Table 1). Of these, 3 samples were excluded due to low alignment rates. Using the 14 remaining samples, we randomly generated a set of 30 “3 versus 3 comparisons” (Supplementary Table 2). For each of the 30 comparisons, the consensus peak set and the normalisation method implementation incorporated only the 6 samples within that comparison.

### Inflation of binding sites and normalisation implementation in simulation experiments

In order to simulate sites of “true” differential binding, we developed a process to artificially inflate peaks in the CTCF CUT&RUN data set. To inflate a peak, the raw counts (reads overlapping with that peak) in each sample within a group in the comparison were multiplied by a factor of 2, 5, 10 or 20. The peaks were randomly selected (without replacement) and the fraction of total peaks that were inflated for an experiment is found in the Results section.

After peak inflation, scaling factors were recalculated for the library size, background, reads-in-peak and spike-in normalisation methods. For library size normalisation, the scaling factors incorporated the additional imputed counts from the entire sample. For background normalisation, imputed reads were added to overlapping background bins, and DiffBind recalculated the scaling factors based on these updated bin counts. For spike-in normalisation, the spike-in BAM counts were scaled in proportion to the increased library size after inflation and spike-in scale factor was recalculated based on this. As the scaling factors for the reads-in-peak method are derived from the peakset itself, they were automatically calculated within DiffBind. For Greenlist, the scaling factors were not recalculated as they are derived from regions of “constant nonspecific signal”, meaning that these regions should be minimally affected by imputing signal.

## Results

### Benchmarking normalisation options

To begin we benchmark five different normalisation options using a variety of synthetic approaches with expected ground truth but with a basis in real experimental data to make our simulations as realistic as possible. We used experimental data from the CTCF CUT&RUN dataset published in (3). Specifically, we designed a series of three experiments to simulate a variety of plausible protein-DNA interacting scenarios using 30 three-versus-three comparisons (runs) described in Supplementary Table 2.

#### Experiment 1: Control of Type I error

As the binding in the CTCF CUT&RUN data set is of one protein with no experimental perturbation, we assume that no statistically significant biological differences between samples should be detected. Therefore, any detected differential binding is due to unwanted variation such as technical and experimental variation and are deemed false positives. A good normalisation approach should hold its size below a 5% false positives. Comparing the differential binding across the combinations of samples from this experiment allows us to evaluate the number of false positives (Type1 error).

Running the analysis for each of the 30 runs of 3 vs 3 samples revealed that the total number of peaks in the consensus peak set varied between runs of different sample combinations from approximately 52,000 to 66,000 peaks. The percentage of peaks found to be significantly differentially bound for each of the 30 runs are presented in Figure 1A. The majority of the runs produce an acceptable number (<5%) of false discoveries across all five normalisation methods. However, spike-in normalisation failed to hold its size in 6/30 (20%) of the runs, and as many as 95% of detected peaks were identified as differentially changed. reads-in-peak and Greenlist failed to hold their size in two runs, library size in one run, and background held its size under 5% false discoveries for all runs. With Run 17, all normalisation approaches performed the worst and were unable to fully account for the technical effects between samples. This may be explained by Run 17’s make up, in which samples in one group were consistently digested at lower average temperatures and received lower sequencing depth (Supplementary Tables 1 and 2).

**Figure 1:**
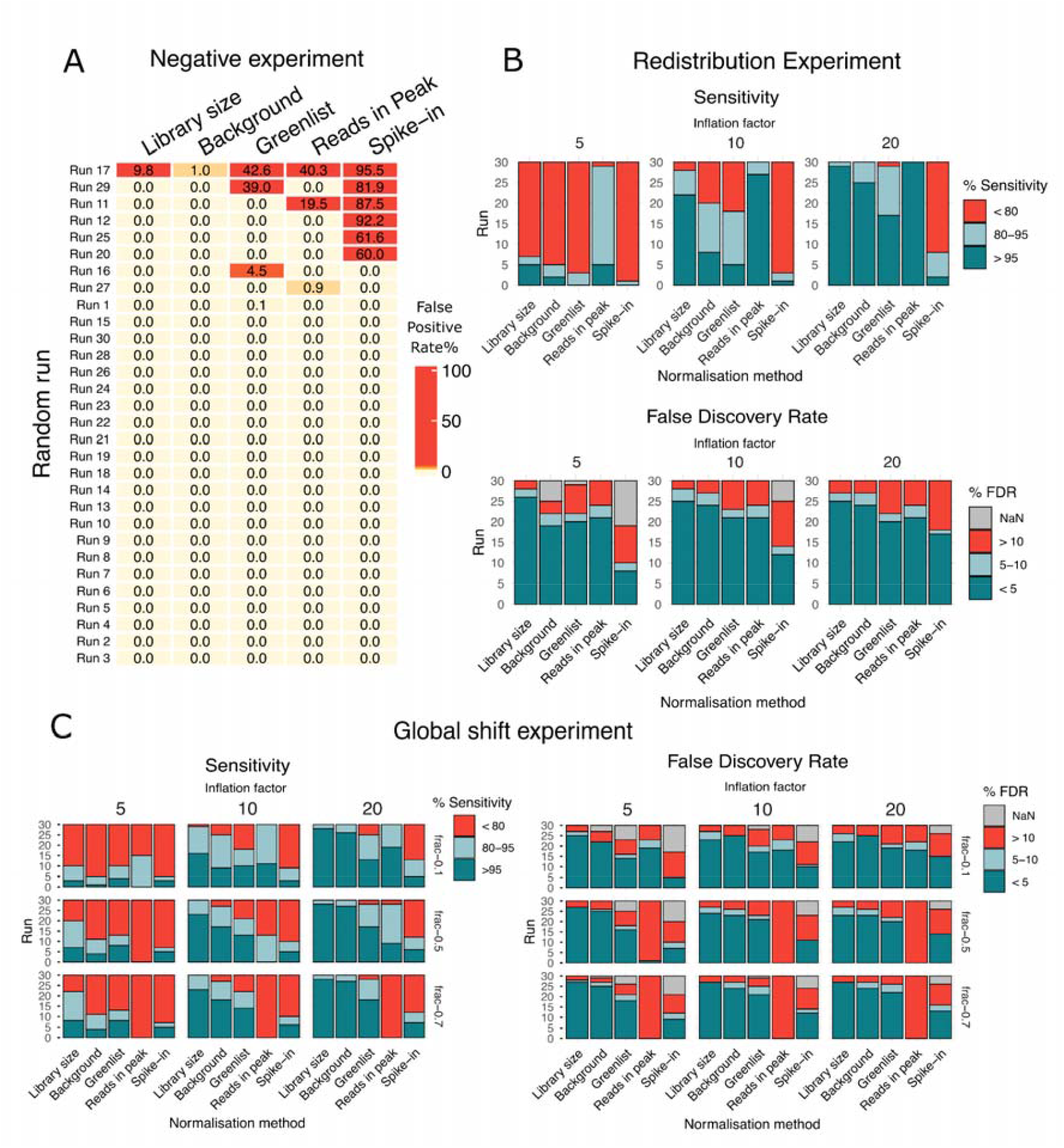
Results of synthetic experiments on CTCF CUT&RUN pooled samples benchmarking the five normalisation methods. A) Negative experiment (Experiment 1): Control for Type 1 error or “Negative Experiment” results. The number of false positives detected by each normalisation method tested is shown in the heatmap for each of the 30 runs. Runs with over 5% of falsely detected differentially bound peaks (as a percentage of the total number of peaks in the consensus peak set for that run) are coloured in red. B) Redistribution experiment (Experiment 2): Sensitivity and False discovery rate of peaks across library size, background, greenlist, reads-in-peak and spike-in normalisation methods with simulated symmetrical or “redistribution of protein binding”. The simulated “true” signal was imputed by a factor of 5, 10 and 20. The bars indicate the number of runs in which the analysis holds its size. Red indicates runs with poor performance while blue is acceptable. C) Similar to B by simulating a global shift (Experiment 3): Sensitivity and False discovery rate of peaks across library size, background, greenlist, reads-in-peak and spike-in normalisation methods with simulated “global shift in protein binding” with inflation factors of 5, 10 and 20.

In summary, background, library size and reads-in-peak normalisation methods were best performing in their ability to minimise technical variation between groups resulting in the least number of regions falsely detected as having increased protein binding.

#### Experiment 2: Investigating a “redistribution” of protein binding

In Experiment 2, we tested the ability of the normalisation methods to detect “true” differences in binding by inserting known changes into the data. For each comparison (Supplementary Table 2), we mimic a “redistribution” of the protein where in one group, the raw counts for 5% of peaks are multiplied or “inflated” by a factor and a different random set of 5% peaks are inflated by the same factor in the samples of the other group. The inflation factors used were 5, 10 and 20 resulting in 90 independent simulations (30 sample combinations x 3 factors of inflation).

We use the metrics of “sensitivity” and “false discovery rate” (FDR) to evaluate the normalisation methods. Sensitivity is the number of true artificially inflated sites that were detected (“true positives”) in the differential analysis as a percentage of all artificially inflated sites regardless of detection (“true positives” + “false negatives”). We consider a sensitivity of >95% as good, a sensitivity between 80 to 95% as acceptable and less than 80% as poor. FDR is the number of differentially bound sites detected that were not inflated (“false positives”) as a percentage of all detected differentially bound sites (“all positives”). We consider an FDR of less than 5% as good, an FDR between 5 to 10% as acceptable and an FDR above 10% as poor. NaN values for FDR indicated that no positives (“true” or “false”) were detected in the differential analysis even after artificially inflating biological signal. The ideal method would maximise sensitivity and minimise the false discovery rate.

The sensitivity of each method increased as the inflation factor increased, as expected, with relatively low sensitivity at 5-fold and below (green colours, Figure 1B and Supplementary Figure 2A - including inflation factor 2). The reads-in-peak normalisation method had the greatest ability to detect differential binding over all inflation factors tested. Library size had acceptable levels of sensitivity for most simulations with larger inflation factors. Background, greenlist and spike-in normalisation methods have lower sensitivity in this context, with spike-in performing the worst.

When considering FDR (Figure 1B and Supplementary Figure 2A), most methods were consistent with respect to the inflation factor. Library size normalisation held its size in more of the simulations than other methods while background and greenlist normalisation methods generally performed well. Reads-in-peak balanced its higher sensitivity with a slightly higher FDR. Of all methods, the spike-in normalisation method had the highest FDR.

In summary, the reads-in-peak normalisation method was the most robust in the context of balanced numbers of up-regulated and down-regulated peaks.

#### Experiment 3: Investigating a “global shift” in protein binding

In this experiment we simulate a scenario where binding globally increases or decreases in response to a condition, mimicking a “global shift” in protein binding. This is plausible in contexts such as gene knockouts (where mutation, silencing, or removal reduces protein expression and binding) or drug activation (where enhancing an existing protein increases its chromatin binding).

In this data set we use the same 30 run sample combinations (Supplementary Table 2) and vary both the proportion of total binding sites with increased binding (0.1, 0.5 and 0.7) as well as the inflation factor by which the amount of binding within a peak occurs at these sites (5, 10 and 20) leading to 270 datasets to analyse for differential binding with each of the normalisation methods (Figure 1C). Further details of each run (including inflation factor 2) are given in Supplementary Figures 2B-D.

The overall sensitivity of the library size and background methods increased with increasing inflation factor and to some extent with a greater fraction of peaks inflated. The reads-in-peak normalisation method behaved differently. When 10% of the peaks were inflated by a factor of 5 or 10, the reads-in-peak normalisation had the greatest sensitivity of all methods tested. However, when 50% or 70% of the total number of peaks were inflated in one group, the assumption of the majority of sites not being changed or the up and down regulation of peaks being symmetric is violated. Therefore the ability of the reads-in-peaks normalisation method to detect the inflated peaks diminished noticeably and in almost all runs, there was low sensitivity and an unacceptable percentage of significant peaks detected were false positives (Figure 1C). These results demonstrate the limited application of the reads-in-peak normalisation method as a viable option in the context of “global” shifts or unidirectional binding profiles. The spike-in normalisation method increased performance with respect to sensitivity and FDR with increasing inflation factors and fraction of peaks inflated, but overall performed the more poorly compared to library size, background and greenlist normalisation methods.

When considering greater inflation factors (10 and 20) and an inflation of larger fraction (0.5 and 0.7) of total peaks, library size and background normalisation methods were comparable in both sensitivity and FDR metrics, with the greenlist normalisation method falling slightly behind these methods. While library size and background normalisation methods resulted in very similar FDR metrics, library size having higher sensitivity overall suggests that in the context of global shift, it is the optimal method.

### Demonstration of global shift capture in MCF7 ER dataset

To evaluate differential binding in a real global shift experiment, we performed CUT&RUN for the estrogen receptor (ER) protein in the ER+ breast cancer cell line MCF7. This experiment included whole-cell drosophila (dm6) and yeast DNA fragment (sacCer3) spike-ins. We varied the amount of ER present in the cells by either the disruption of the *ESR1* gene through CRISPR, or by varying the presence of estradiol (E2), a hormone that activates ER and increases ER binding to the chromatin. The four conditions are summarised in Table 2 with n = 2 replicates per condition; (1) charcoal-stripped serum (CSS) supplemented with estradiol (E2), (2) CSS without supplemented estradiol (noE2), (3) complete medium with ER gene not disrupted (sgControl), and (4) complete medium with ER gene disrupted through CRISPR (sgER). In addition to whole-cell and DNA spike-in normalisations, library size, background, reads-in-peak, and greenlist normalisation methods were also evaluated.

**Table 1:** Summary of normalisation methods, the technical biases they account for and their limitations. Background bins refer to the genomic bins generated by csaw (wrapped in DiffBind) of 15,000 basepairs. Cleavage efficiency refers to the proportion of fragments collected that are from protein bound regions compared to outside these regions.

| Normalisation method | Principle/ Rationale | Technical variation addressed | Limitations/ Assumptions |
| --- | --- | --- | --- |
| Library size | In sequencing, the number of reads in a | Accounts for sequencing depth | Does not account for many other sources of |
|  | peak is proportional to the total number of fragments collected per sample. | variation between samples | technical variation. |
| Reads in Peak | Assumes each sample has a similar number of bound molecules and the majority of peaks are not differentially bound, or there is balance between up and down regulated peaks. | Accounts for changes in cleavage efficiency between samples. | Assumption of a similar number of bound molecules across conditions may not hold true in global shifts. |
| Background | The underlying assumption is that the majority of background bins are not subject to differential enrichment across samples. | Accounts for systematic fold changes in the coverage of the background bins (25). | Assumes a consistent cleavage efficiency in the CUT&RUN assay across samples. Some biological perturbations may violate this assumption. |
| Greenlist | Empirically derived regions that are assumed to have consistent levels of noise (not specific for the protein) in all samples. | Accounts for variation in cleavage efficiency and sequencing depth variation between samples. | The regions may not meet the assumptions in unusual conditions. This method is only available for some organisms. |
| Spike-in | A constant amount of spike-in DNA or cells in every sample provides a quantitative reference to which to calibrate reads in downstream analysis. | Accounts for enrichment relative to the number of cells in the sample, sample loss, sequencing depth variation, composition bias | Prone to liquid handling measurement error and antibody cross-reactivity (in whole-cell spike-in) |

**Table 2:** Description of the four conditions in the ER CUT&RUN experiment and a qualitative estimate of expected ER chromatin binding levels based on the biology. The complete medium contains phenol and the fetal bovine serum (FBS) contains “ambient” levels of estrogen, both of which activate ER and increase its chromatin binding capacity. The *ESR1* gene is disrupted in the sgER condition, but not in the sgControl condition. In the E2 and noE2 conditions, Phenol-free medium and charcoal stripped FBS deprived cells of hormones and the ESR1 gene is not disrupted. In the E2 condition, estradiol was added one hour before cells were collected to activate ER, maximising chromatin binding levels so we expect the average amount of ER protein binding per cell to be the greatest out of the four conditions. In the noE2 condition estrogen is stripped from the media, but the *ESR1* gene remains intact, so some amount of ER is expected to be present.

| Condition | sgControl | sgER | E2 | noE2 |
| --- | --- | --- | --- | --- |
| Media | Complete | Complete | Charcoal stripped | Charcoal stripped |
| *ESR1* gene disruption through CRISPR | NO | YES | NO | NO |
| Estradiol supplementation | NO | NO | YES | NO |
| Expected overall ER binding level | <u>Moderate</u>; *ESR1* gene and ER protein undisrupted, estrogens are present, but ER turnover rate is high. | <u>Low</u>; *ESR1* gene and ER protein disrupted, E2 present at moderate amounts. | <u>High</u>; *ESR1* gene and ER protein undisrupted, ER highly activated by E2 hormone without enough time to degrade. | <u>Low</u>; *ESR1* gene and ER protein undisrupted, but activating E2 hormone amount is low. |

Each sample was sequenced with a depth of around 30M reads (Supplementary Table 3) and 200-1800 peaks were detected depending on the sample set using MAC2 (24). The read enrichment heatmaps of the unnormalised samples is shown in Supplementary Figure 4B. The number of peaks called by condition broadly agrees with the expected biology, with the E2 stimulation having on average the highest and the sgER condition having the lowest (Supplementary Table 3).

Principal Component Analysis (PCA) was performed on the unnormalised (Supplementary Figure 5) and normalised samples (Supplementary figure 6A-F) using the scaling factors derived from each normalisation method to illustrate sample clustering. The PCA plots suggested that spike-in normalisation methods increased the distance between replicates within a condition. To quantify replicate concordance, we calculated the Euclidean distance between replicates of the same condition. As the purpose of normalisation is to minimise variation due to technical effects, a reliable normalisation method should improve replicate concordance within samples of the same experimental condition while preserving true differences between the biological factors of interest. As shown in Figure 2A, library size and background normalisation consistently reduced intra-replicate variability relative to unnormalised counts. Greenlist and reads-in-peak methods were less consistent but generally reduced distance between replicates. In contrast, both spike-in methods (sacCer3 DNA and dm6 whole-cell) dramatically increased intra-replicate variability for some conditions. We speculate that the reason for the increased Euclidean distance in the sgER and E2 groups when sacCer3 spike-in is implemented is due to noticeably different scale factors resulting from the large variability in yeast aligned reads. Likewise, for the dm6 whole-cell spike-in, the decrease in replicate concordance of the sgControl group is explained by the variable number of dm6 reads. The decrease in replicate concordance of the sgER group with dm6 whole-cell spike-in is slightly harder to explain but may be the result of the slight difference in sequencing depth between the two samples.

**Figure 2:**
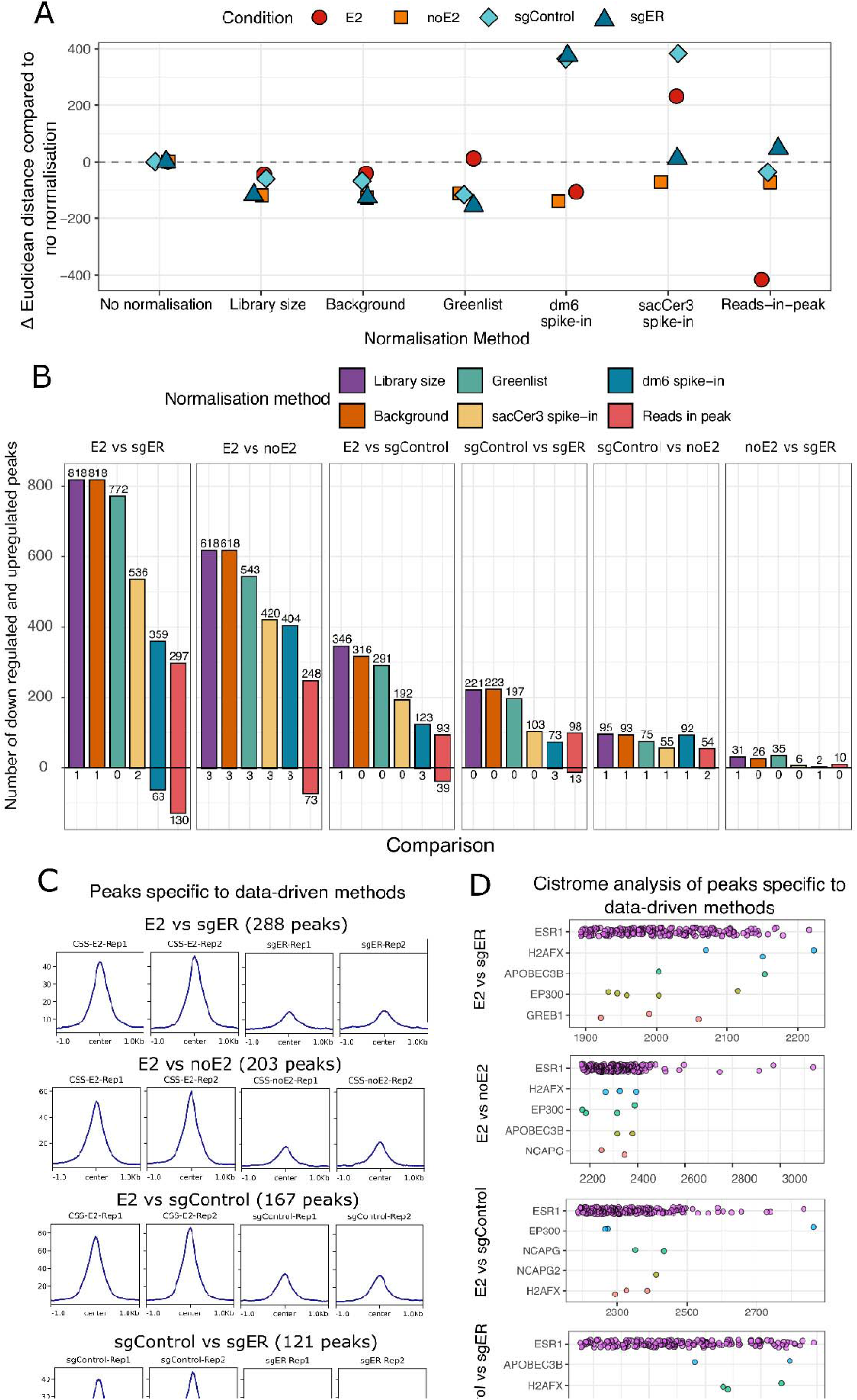
Analysis of ER CUT&RUN dataset on MCF7 cells. A: Difference in Euclidean distance of the normalised values between each replicate compared to non-normalised values within a condition by normalisation method. No normalisation is also shown. B: Pairwise comparisons of all conditions described in Table 2. Statistically significant (FDR<0.05) up-regulated peaks are above the x-axis and down-regulated peaks are below the x-axis. C: Raw unnormalised aggregated signal of peaks in each comparison that are data-driven specific (i.e. found in library or background normalisation methods but not in the spike-in methods) in E2 versus sgER, E2 versus noE2, E2 vs sgControl and sgControl versus sgER comparisons. Read enrichment plots for this analysis with all comparisons are found in Supplementary Figure 7. D: Cistrome analysis of peaksets in Figure 2C, showing top 5 factors that have the highest similarity as measured by the GIGGLE score. The GIGGLE score quantifies the similarity using -log10(pvalue) x -log2(odds ratio)(30). Each dot represents a sample from a publicly available dataset (transcription factor/chromatin regulator). Cistrome analysis for all comparisons are in Supplementary Figure 8.

Differential binding analysis was performed across all pairwise conditions, yielding six comparisons in total. Two of these comparisons had clear, biologically grounded hypotheses: E2 vs. noE2 was expected to produce many up-regulated peaks with few to no down-regulated peaks, while sgControl vs. sgER was expected to yield exclusively up-regulated peaks with none down-regulated. The remaining four comparisons, where both the expected estradiol level and ER protein abundance vary simultaneously, are slightly more complicated to predict.

The number of differentially bound peaks of all six pairwise comparisons are presented in Figure 2B. For comparisons where a global shift in binding was anticipated, we reasoned that the normalisation method identifying the greatest number of up-regulated differentially bound peaks would reflect superior sensitivity, while unexpected down-regulated peaks would be indicative of false positives.

Figure 2B shows that across nearly all comparisons, library size and background normalisation identified the greatest number of differentially bound up-regulated peaks, indicating the greatest sensitivity of the normalisation methods tested. For the comparisons where the biology is well characterised (E2 versus no E2 and sgControl versus sgER), library size and background normalisation methods identify roughly double the number of up-regulated peaks compared to the spike-in methods. The whole-cell spike-in method identified 3 down-regulated peaks in the sgControl versus sgER comparison, which given the hypothesis of exclusively upregulated peaks, are false positives.

In the remaining comparisons the biology is not as well characterised because the comparisons vary across media as well as presence of estradiol and ER protein abundance. However, we can infer expected directions of regulation across conditions as reasoned in Table 2. For example, in the E2 versus sgER comparison, we expect there will be a trend towards an upregulation of peaks as there will be an overall large amount of ER binding to the chromatin in the E2 condition and a lower amount of ER binding in the sgER condition where the gene has been disrupted. In this comparison, a greater number of differentially bound peaks are found in the library size and background methods (818 up-regulated and 1 down-regulated peak) compared to the spike-in methods which detect approximately half or fewer upregulated peaks.

We next wished to verify that the extra differential peaks identified by the library size and background normalisation methods represent genuine ER binding sites and not false positives. Therefore, we derived a “data-driven-specific” peak set by taking the union of peaks found to be differentially bound from the library size and background normalisation methods and subtracting the union of differentially bound peaks from the sacCer3 and dm6 spike-in methods. The strong binding enrichment signal in data driven peaks (Figure 2C) gives evidence that the differentially bound peaks are true positives. This peak set was also compared with publicly available ChIP-seq datasets using the CistromeDB Toolkit(31). Indeed we found strong similarity of the data driven peak with publicly available datasets where ER binding was profiled on the chromatin, i.e. ESR1 (Figure 2D). This provides evidence that the unique peaks identified by library size and background normalisation approaches represent true ER binding events rather than false positives. The read enrichment heatmaps and cistrome analysis for the data-driven peaks for all comparisons are found in Supplementary Figures 7 and 8.

Taken together, the library size and background normalisation demonstrated the strongest overall performance, with greenlist normalisation performing comparably but with marginally reduced sensitivity. In contrast, whole-cell Drosophila and yeast DNA spike-in normalisations captured fewer up-regulated peaks across the majority of comparisons relative to library size and background normalisation. Likewise, reads-in-peaks normalisation produced a higher proportion of down-regulated peaks across comparisons, consistent with the inherent assumptions of this approach and suggestive of an elevated false-positive rate in this context.

### Different normalisation methods results in discrepant interpretations in MCF7 BRG1 dataset

To highlight the extent to which normalisation choice can result in discrepant biological interpretation and not just differences in sensitivity, we analysed CUT&RUN for the BRG1 protein on MCF7 cells treated with DMSO or abemaciclib. Abemaciclib is a cyclin-dependent kinase 4 and 6 inhibitor, which induces cell-cycle arrest in these cells (32). BRG1, a subunit of the SWI/SNF chromatin remodelling complex, is canonically involved in nucleosome repositioning and hence changes in local chromatin accessibility. Yeast DNA *(Saccharomyces cerevisiae)* was included for the implementation of spike-in normalisation as described in the methods section. The number of reads aligned to the concatenated hg38–sacCer3 genome and to each individual species (after removing reads overlapping blacklisted regions), along with fraction of reads in peaks (FRiP) scores and MACS2 peak counts, are summarised in Supplementary Table 4. The raw unnormalised peak signal of each sample is visualised in Supplementary Figure 9E.

PCA plots of the samples under different normalisation methods are given in Supplementary Figure 9A. Reassuringly, in all normalisation methods, PC1 is explained by treatment condition and PC2 follows replicate variation and methods could both reduce and increase the difference between replicates (Supplementary Figure 9B).

After testing for differential binding the resulting number of significantly differentially bound peaks (FDR < 0.05) varied substantially between normalisation methods (Figure 3A). For example, background, greenlist and library size normalisation called less than 140 peak regions with increased BRG1 binding (up-peaks) and many more down regulated peaks. In contrast, the spike-in normalisation resulted in fewer significantly changed peaks overall but had the highest number of up-peaks. Strikingly, the agreement or intersection between all methods is less than 11% of the union of significant peaks (234/2287) (Figure 3B). To confirm that this is not the result of an arbitrary FDR threshold we compared the FDR values and ranks between the different normalisation methods, and indeed, the same peaks have vastly different FDR values and ranking depending on the normalisation method (Supplementary Figure 9C & 9D).

**Figure 3:**
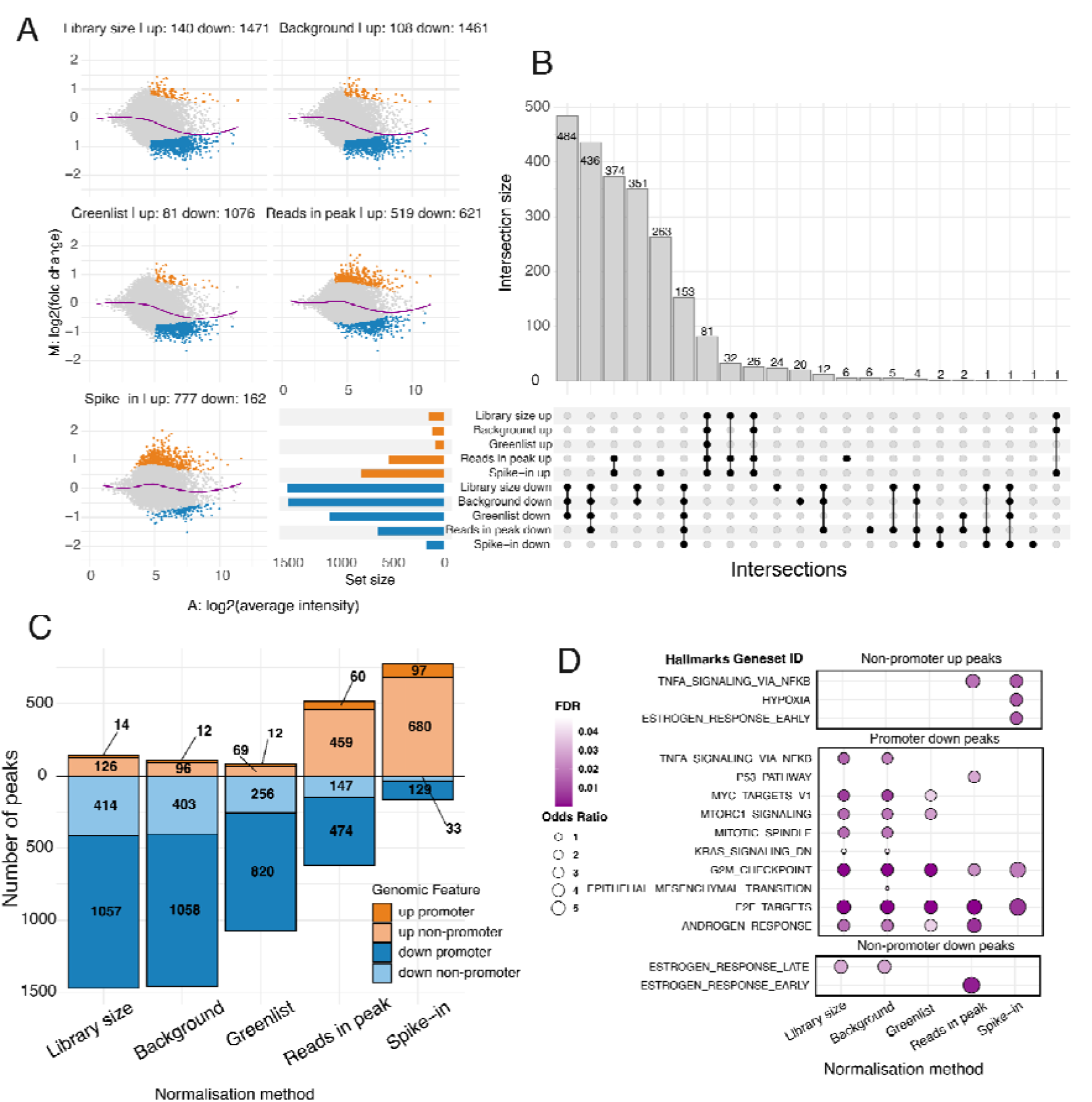
Differential analysis with five normalisation methods applied to the BRG1 CUT&RUN dataset. Here, “up” refers to increased protein binding in the treatment condition, abemaciclib, when compared to the control condition DMSO. A) MA plot visualisations of the same data normalised by 5 different methods: library size, background, reads-in-peak, greenlist and spike-in methods with the number of differentially up and down peaks resulting from each normalisation method annotated in the title. B) Upset plot showing common and unique sets of significantly differentially bound peaks across the five normalisation methods. C) Direction and genomic location breakdown (by promoter versus non-promoter) of significantly differentially bound peaks by normalisation method. D) Hallmarks of Cancer gene set enrichment of peaks stratified by direction and genomic location, computed using ChIP-Enrich. Peaks were assigned to genes by the nearest transcription start site. Odds ratios reflect the association between peak presence and gene set membership, controlling for gene locus length.

We categorized differentially bound peaks by genomic location (promoter versus non-promoter), creating four groups: up-promoter, up-non-promoter, down-promoter, and down-non-promoter. The distribution varied by normalization method with background, greenlist, and library size methods having a majority of down-regulated peaks at promoters, while reads-in-peak and spike-in normalization methods resulted in a majority of up-regulated peaks at non-promoter regions (Figure 3C).

To characterise the differences in biological interpretation of BRG1 chromatin binding using different normalisation methods, we performed gene set enrichment analyses on the genes associated with each of these four groups of peaks (Figure 3D). The cell-cycle arrest gene sets (G2M_CHECKPOINT and E2F_TARGETS) were the only two gene sets commonly enriched in down-regulated promoter peaks across all five normalisation methods tested. Strikingly, 11 gene sets were enriched in only some normalisation methods, demonstrating that normalisation choice can substantially influence pathway enrichment results and potentially direct researchers toward different experimental directions.

We further analysed the total BRG1 levels in the nucleus with DMSO and abemaciclib *500 nM for 2 days* with a western blot (Supplementary Figure 9G). The western blot showed a slight increase in total nuclear (sum of bound and unbound) levels of BRG1 protein in the abemaciclib condition. The spike-in normalisation method is the only method that follows the trend of increased BRG1 binding to the chromatin. However, we do not see substantial changes to the number of peaks called between conditions by MACS2 (Supplementary table 4) and the proportion of peaks identified as differentially bound is small relative to the total number in the consensus peakset (21,880 peaks). Therefore, in this case with vastly different results for different normalisation methods validation would need to be performed at the individual peak level.

Overall the analysis of this data set demonstrates that the choice of normalisation method significantly impacts differential binding analysis results, not only with respect to the numbers of differentially bound peaks, but also the downstream biological interpretation. Secondly, when this case occurs, an orthogonal method of probing the system is required (e.g. the qPCR of specific regions).

## Discussion

There are currently no clear guidelines for the optimal normalisation method to use in the context of differential binding analysis in CUT&RUN. This is a significant issue for reproducibility of analysis and the reliability of downstream interpretation of data. We demonstrate this issue with in-house generated datasets and public data where we conduct a series of simulations to benchmark five normalisation methods. Of the methods we considered, library size and background normalisation were the most robust in a variety of plausible binding scenarios.

To systematically test which normalisation methods would perform the best in a variety of plausible CUT&RUN differential analysis scenarios, we conducted a series of experiments. In synthetic experiment 1 we evaluated the detection of false positives in the CTCF dataset. As these samples were not biologically perturbed and so the true amount of protein binding at the chromatin is not expected to vary, we expected no differentially bound peaks. In this experiment spike-in normalisation produced the greatest number of false-positives. To determine which normalisation methods were able to detect “true signal”, we imputed signal into this dataset in synthetic experiment 2 and 3. The reads-in-peak method performed the best in the context of protein redistribution in synthetic experiment 2, with library size normalisation also performing well. Here, 10% of total binding sites within a run were artificially inflated, leaving the majority of binding sites unchanged across conditions. The reads-in-peak method was originally created in analogy to RNA-Seq where it is assumed the majority of genes (or in our case binding sites) are not expected to be differentially expressed or have symmetric up and down regulation. Experiment 2, protein redistribution, is aligned with this assumption and so it is perhaps not surprising that reads-in-peaks performed the best in this experiment.

In synthetic experiment 3, we test the normalisation methods in the context of global shift of protein binding across conditions, plausible in the case of gene knockout or protein-activating drugs. Library size and background normalisation methods were the best performing in this context.

In the ER CUT&RUN dataset we examined all normalisation methods tested in the simulation experiments, including whole-cell and DNA spike-in. The findings of the global shift simulations are broadly recapitulated in the real ER dataset. When examining replicate concordance using Euclidean distance as a metric, library size, background and greenlist normalisation methods increased replicate concordance within all conditions, whereas the spike-in methods and reads-in-peak decreased it. A method that increases inter-replicate distance after normalisation when compared to raw counts will likely be a less sensitive method. In our synthetic and ER datasets, we demonstrated the unreliability of spike-in normalisation in the context of differential binding analysis.

In the BRG1 dataset, the choice of normalisation method substantially altered the number, direction and genomic location of differentially bound regions, and led to markedly different enrichment results. Here, we highlight that where the biology is unknown, an orthogonal method of assessing binding behaviour may be required to validate the choice of normalisation method. We speculate that in this system the use of abemaciclib may result in the production of non-specific DNA fragments (i.e. the biological perturbation itself causes a change in the efficiency of the CUT&RUN assay), which is unlikely to be the case with the biological perturbations the ER dataset. This is apparent in Supplementary Figure 3B where after spike-in normalisation, the abemaciclib condition has increased inter-replicate distance.

There are two main types of spike-in normalisation; with exogenous naked DNA fragments or chromatin and with whole cells. DNA fragment spike-in normalisation is subject to strict assumptions such as the spike-in fragments, the protein target and the corresponding antibody behave in the same manner as their endogenous counterparts throughout the protocol. It also assumes the ratio of spike-in chromatin to endogenous chromatin is the same across all samples. These assumptions are hard to verify in a practical setting. Being a method reliant on wet-lab protocol, it is susceptible to errors such as hand-to-hand error, and antibody cross-reactivity (in the case of whole-cell spike-ins). Finally, there isn’t a consensus on the most robust and reproducible protocol for implementing spike-in normalisation in the analysis pipeline(8). For these reasons we discourage spike-in normalisation in the context of CUT&RUN differential analysis where we wish to compare binding between conditions.

CUT&RUN uses Micrococcal Nuclease (MNase) tethered to specific antibodies to cleave DNA flanking target protein epitopes, producing lower noise than ChIP-Seq’s sonication-based fragmentation. The protocol further reduces background noise by filtering read fragments >120bp, justified by periodicity in fragment length distributions indicating nucleosomal cutting. Background normalization is discouraged in the CUT&RUN protocol due to low inherent noise(3). However, we did not apply the 120bp filter as it removes a substantial number of reads and therefore signal, particularly for proteins with fewer binding sites. In our experience, not all proteins show fragment length periodicity, possibly due to larger complex formation or indirect DNA binding.

The CTCF CUT&RUN samples used in our synthetic experiments varied in digestion time, temperature, cell number, and sequencing depth (Supplementary table 1). Since digestion temperatures above 0°C increase background noise, samples with suboptimal conditions may violate the assumption of biologically unchanging genomic bins required for background normalization. Additionally, the samples varied greatly in their read counts. We suspect this explains why library size normalization generally outperformed background normalisation in our analysis, despite the latter’s rationale aligning well with CUT&RUN differential analysis. In our synthetic experiments, we scaled the number of spike-in reads for a sample based on the increased library size that was caused by inflation. This approach to simulating spike-in normalisation factors has associated limitations as spike-in are inherently a wet-lab method. Specifically, any errors with spike-in implementation in the original dataset would be carried through in synthetic experiments 2 and 3.

Greenlist normalisation derived a set of genomic regions for the human genome that have consistently “high entropy” or “constant nonspecific signal” based on the analysis of 463 publicly available human IgG CUT&RUN samples(10). This method did not perform as well as library size and background normalisation methods in our experiments but had stable performance across experimental settings. As mentioned by the authors, this method requires the presence of a substantial number of already-published IgG samples for the greenlist regions to be determined for a new species, currently limiting its usability to human and mouse species.

In this study, we focussed on protein interactions with the DNA rather than histone modifications in CUT&RUN assays. Histone modifications are much more widely distributed across the genome and experience with ChIP-seq analysis indicated that these analyses require different approaches across the analysis pipeline from peak identification through to comparisons between samples(33–35).The conclusions drawn in our analysis are unlikely to apply directly to differential binding analysis of histone modifications. Indeed, there have been examples where spike-ins have uncovered the expected biology whereas data-driven methods have overnormalised(36). However there exist data-driven methods to analyse histone modifications where global shifts may be expected though window-based methods such as ChIPseqSpikeInFree(7) and csaw(25) which may enhance reproducibility of results.

Our investigations were limited to five normalisation methods and peak-based methods (as opposed to window-based). While other methods of normalisation are available, we did not test them as they are unlikely to be suitable for CUT&RUN differential analysis because of the assumptions they make about the data. The choice of peak-based or window-based approaches in a transcription factor depend on expected binding behaviour.

The choice of normalisation for a particular dataset requires some investigation or prior knowledge of the biological system. Considering the stochastic nature of spike-ins (i.e. while they may excel in some cases, they can also fail imperceptibly), we offer a set of recommendations of which normalisation method to implement in the context of differential binding analysis of transcription factors in CUT&RUN based on the results of the simulation studies. As always, the best methods are those in which the data best match the assumptions of the normalisation approach. For each normalisation method tested in this study, these are the assumptions for which the methods perform well:

1. Reads-in-peak normalisation assumes that either most peaks are not changing their binding profile or they are balanced in up/down regulation. When these assumptions are satisfied it is the best performing method. However, it is rarely known for certain that these assumptions will be met.
2. When a “global shift” in binding profiles is suspected, we recommend the implementation of the library size normalisation, which is the default in DiffBind. Background normalisation also performs well in the context of global shifts. However, it may be more susceptible to differences in overall distribution of a protein with very promiscuous binding, violating the assumptions that most bins without detected peaks only contain background reads.
3. If the behaviour of the protein of interest is unknown, we recommend the use of the default library size normalisation. This is a conservative approach and will minimise the likelihood of non-reproducible downstream interpretations.
4. There are certain biological systems that might not be able to be analysed using data-driven methods such as when the biological perturbation causes a large technical variation as well (e.g. DNA damage). In this case, we suggest that the use of spike-ins should be validated by an orthogonal assay to provide ground truth.

## Conclusion

In the context of differential binding analysis, we find that of the normalisation methods tested, data-driven methods are the most robust methods in a variety of contexts including simulated global shifts and a redistribution of protein binding across experimental conditions. Spike-ins which purport to account for global shifts are subject to wet-lab variation, antibody cross-reactivity as well as unclear guidelines of implementation. We suggest their use should be justified with orthogonal assays.

## Author Contributions

**K. Ambani:** Data curation, formal analysis, investigation, writing–original draft, writing-review and editing. **A. Ahn:** Formal analysis, writing–review and editing. **A.C. Watt:** writing–original draft, writing-review and editing, investigation. **C. Blyth:** Investigation. ***B. Russ:*** Investigation. **M. Taylor:** Investigation. **S. Goel:** Resources, supervision, funding acquisition, writing–review and editing. **A. Oshlack:** Conceptualisation, data curation, supervision, writing–original draft, writing–review and editing.

## Data Availability

The sequencing data for the BRG1 CUT&RUN experiment is publicly available in Gene Expression Omnibus (GEO), accession number GSE331453. The ER CUT&RUN data set is available at GEO Accession GSE331454. The CTCF CUT&RUN dataset is from the original CUT&RUN protocol paper(3), GEO Accession number GSE84474. The code that processed the raw files, conducted the differential analysis on the BRG1 and ER experiments, simulation experiments for the Henikoff datasets and the code for figure generation are available here https://github.com/k-ambani/CUT-RUN_normalisation.

