## Supplementary Material for "Benchmarking normalisation methods for differential binding analysis in CUT&RUN"

Supplementary Figure 1: CTCF samples raw signal read-enrichment heatmap	2

Supplementary Table 1: Pool of CTCF CUT&RUN samples	3

Supplementary Table 2: List of comparisons in the simulations	4

Supplementary Figure 2A: Redistribution experiment - sensitivity and FDR	5

Supplementary Figure 2B: Global shift for 0.1 of peaks - sensitivity and FDR.	7

Supplementary Figure 2C: Global shift for 0.5 of peaks - sensitivity and FDR.	9

Supplementary Figure 2D: Global shift for 0.7 of peaks - sensitivity and FDR.	11

Supplementary Figure 3A: ER dataset - Insert size distribution	13

Supplementary Figure 3B: ER dataset - raw signal read-enrichment heatmap	13

Supplementary Table 3: ER dataset - aligned reads, spike-in scale factor and peak counts.	14

Supplementary Figure 4: Western Blot validating ER knockdown	14

Supplementary Figure 5: PCA plot of ER datasets before normalisation	15

Supplementary Figure 6: PCA plot of ER datasets with different normalisation methods	16

Supplementary Figure 7: Read-enrichment heatmaps of significant peaks unique to library size and background methods, not found using spike-in methods	17

Supplementary Figure 8: Cistrome of peaks unique to library size and background methods, not found in spike-in methods	18

Supplementary Table 4: BRG1 - aligned reads, peak numbers and fraction of reads in peak	19

Supplementary Figure 9: BRG1 dataset Quality Check	20

Supplementary Figure 9F: BRG1 dataset insert size distribution	21

Supplementary Figure 9G: BRG1 western blot	21

### Supplementary Figure 1: CTCF samples raw signal read-enrichment heatmap

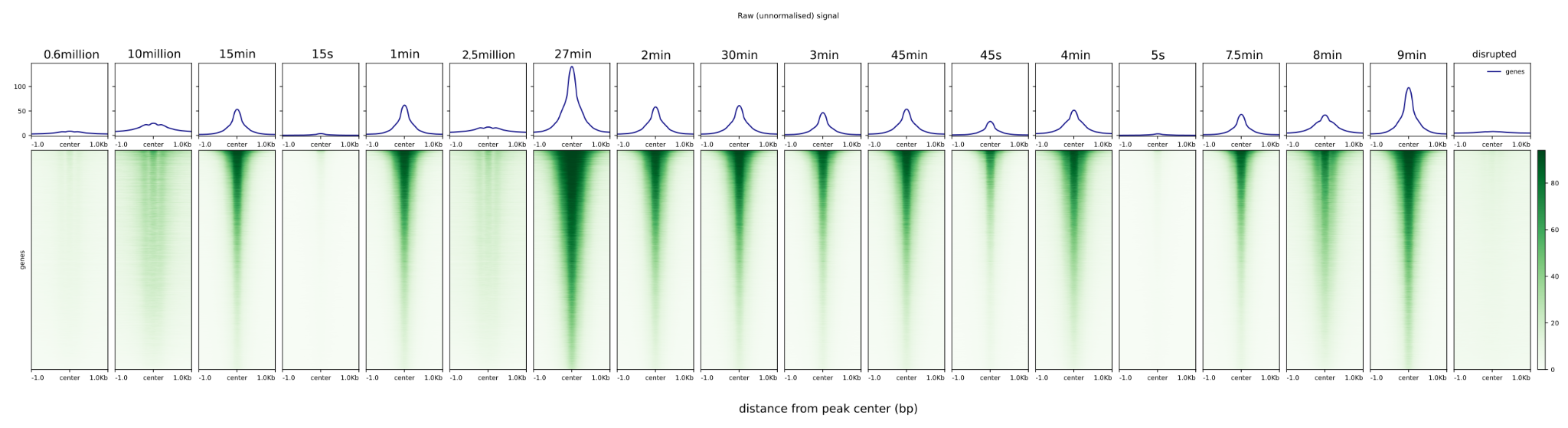
Samples 5s, 15s, 45s and disrupted were excluded from the benchmarking experiments as they had low alignment rates.

### Supplementary Table 1: Pool of CTCF CUT&RUN samples

| **Sample name** | **Sample description from GEO** | **Number of cells** | **Digestion time** | **Digestion temperature** | **Final hg38 bam counts** | **Peak numbers** |
| --- | --- | --- | --- | --- | --- | --- |
| 0.6mill | Cut-and-Run CTCF 0.6 million cells @37C | 0.6mill | ? | 37C | 14,241,624 | 5,792 |
| 2.5mill | Cut-and-Run CTCF 2.5 million cells @37C | 2.5mill | ? | 37C | 28,356,805 | 22,866 |
| 10mill | Cut-and-Run CTCF 10 million cells @37C | 10mill | ? | 37C | 27,943,269 | 41,090 |
| excluded* | Cut-and-Run CTCF 5 s digestion time on ice | 10mill | 5s | 0C | 2,294,986 | 3,159 |
| excluded* | Cut-and-Run CTCF 15 s digestion time on ice | 10mill | 15s | 0C | 2,989,991 | 3,748 |
| excluded* | Cut-and-Run CTCF 45 s digestion time on ice | 10mill | 45s | 0C | 6,925,741 | 32,034 |
| 3min | Cut-and-Run CTCF 3 min digestion time on ice | 10mill | 3min | 0C | 9,146,414 | 45,516 |
| 7.5min | Cut-and-Run CTCF 7.5 min digestion time on ice | 10mill | 7.5min | 0C | 10,277,681 | 37,786 |
| 9min | Cut-and-Run CTCF 9 min digestion time on ice | 10mill | 9min | 0C | 17,249,610 | 59,143 |
| 15min | Cut-and-Run CTCF 15 min digestion time on ice | 10mill | 15min | 0C | 12,090,934 | 45,626 |
| 27min | Cut-and-Run CTCF 27 min digestion time on ice | 10mill | 27min | 0C | 29,128,917 | 75,142 |
| 30min | Cut-and-Run CTCF 30 min digestion time on ice | 10mill | 30min | 0C | 14,282,237 | 52,885 |
| 45min | Cut-and-Run CTCF 45 min digestion time on ice | 10mill | 45min | 0C | 13,745,346 | 52,268 |
| 1min | Cut-and-Run CTCF 1 min digestion time at room temp | 10mill | 1min | 25C | 12,916,312 | 52,661 |
| 2min | Cut-and-Run CTCF 2 min digestion time at room temp | 10mill | 2min | 25C | 13,347,996 | 53,545 |
| 4min | Cut-and-Run CTCF 4 min digestion time at room temp | 10mill | 4min | 25C | 15,626,665 | 56,404 |
| 8min | Cut-and-Run CTCF 8 min digestion time at room temp | 10mill | 8min | 25C | 17,611,713 | 52,574 |

*excluded because of low alignment rate.

Overview of samples used in the synthetic experiments. Samples are from optimisation of the CUT&RUN protocol from the Henikoff paper. A variety of technical parameters were varied. Peak numbers are the number of peaks called by MACS2 for that sample.

### Supplementary Table 2: List of comparisons in the simulations

| Run # | Comparison (group B vs group A) | Run # | Comparison (group B vs group A) |
| --- | --- | --- | --- |
| 1 | 2min, 1min, 30min vs 2.5mill, 45min, 7.5min | 16 | 4min, 2.5mill, 9min vs 30min, 7.5min, 1min |
| 2 | 45min, 2min, 8min vs 4min, 30min, 1min | 17 | 4min, 10mill, 8min vs 2min, 7.5min, 3min |
| 3 | 10mill, 7.5min, 8min vs 30min, 9min, 2.5mill | 18 | 3min, 45min, 2min vs 4min, 9min, 2.5mill |
| 4 | 30min, 27min, 10mill vs 2.5mill, 3min, 8min | 19 | 45min, 30min, 4min vs 0.6mill, 8min, 7.5min |
| 5 | 27min, 15min, 4min vs 3min, 7.5min, 30min | 20 | 7.5min, 2min, 15min vs 8min, 3min, 10mill |
| 6 | 2min, 15min, 2.5mill vs 30min, 45min, 4min | 21 | 0.6mill, 27min, 45min vs 7.5min, 8min, 9min |
| 7 | 45min, 1min, 0.6mill vs 2min, 27min, 30min | 22 | 0.6mill, 3min, 2.5mill vs 8min, 7.5min, 30min |
| 8 | 3min, 45min, 2min vs 15min, 27min, 1min | 23 | 2.5mill, 15min, 1min vs 2min, 8min, 10mill |
| 9 | 8min, 7.5min, 2.5mill vs 45min, 0.6mill, 3min | 24 | 3min, 0.6mill, 1min vs 15min, 45min, 9min |
| 10 | 4min, 45min, 1min vs 10mill, 27min, 30min | 25 | 15min, 3min, 27min vs 45min, 30min, 10mill |
| 11 | 1min, 10mill, 0.6mill vs 9min, 3min, 15min | 26 | 0.6mill, 2min, 9min vs 1min, 45min, 7.5min |
| 12 | 10mill, 2.5mill, 4min vs 1min, 27min, 0.6mill | 27 | 27min, 1min, 2min vs 8min, 15min, 2.5mill |
| 13 | 45min, 27min, 15min vs 30min, 2min, 8min | 28 | 7.5min, 8min, 45min vs 10mill, 1min, 30min |
| 14 | 2.5mill, 8min, 1min vs 0.6mill, 45min, 10mill | 29 | 3min, 15min, 7.5min vs 4min, 45min, 9min |
| 15 | 2min, 4min, 15min vs 3min, 2.5mill, 27min | 30 | 2.5mill, 45min, 1min vs 27min, 15min, 10mill |

The list of comparisons randomly generated from the pool of samples given in Supplementary table 2. These comparisons were kept constant for all synthetic experiments. Details by individual run are found in the following Supplementary Figures 3A-D.

### Supplementary Figure 2A: Redistribution experiment - sensitivity and FDR

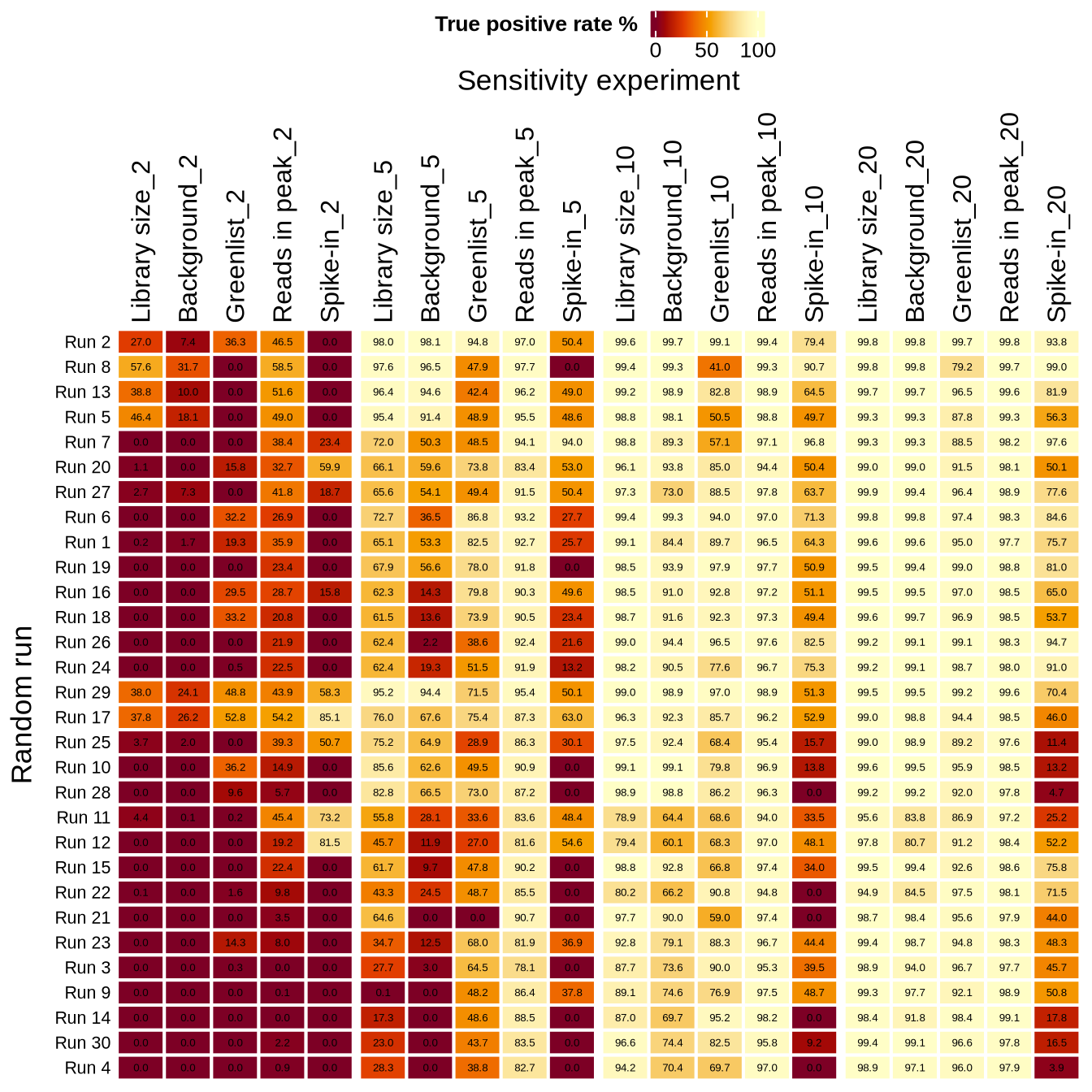

Sensitivity analysis of the redistribution experiment by run. Inflation factors are given in the column names.

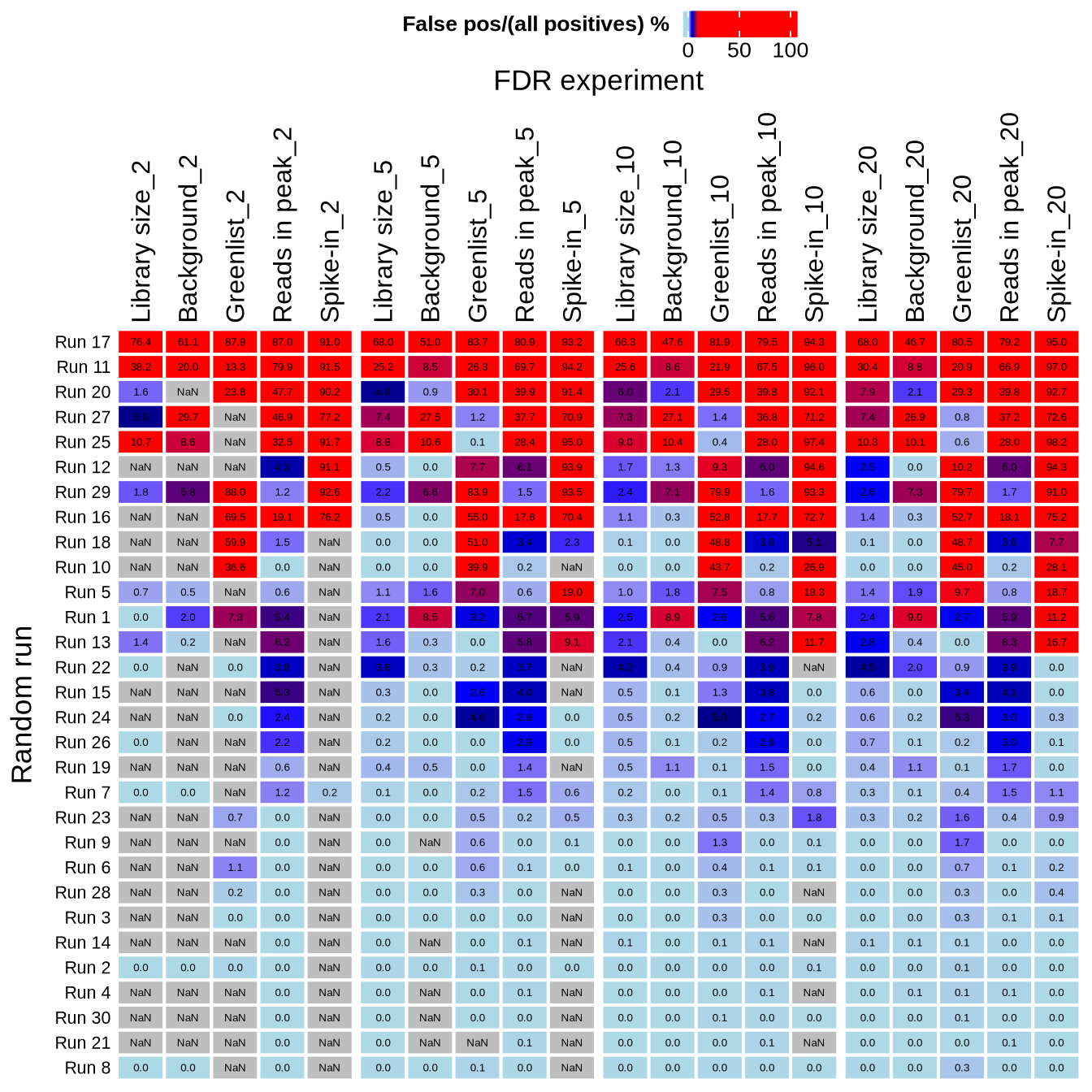

False Discovery rate of the redistribution experiment by run. Inflation factors are given in the column names.

### Supplementary Figure 2B: Global shift for 0.1 of peaks - sensitivity and FDR.

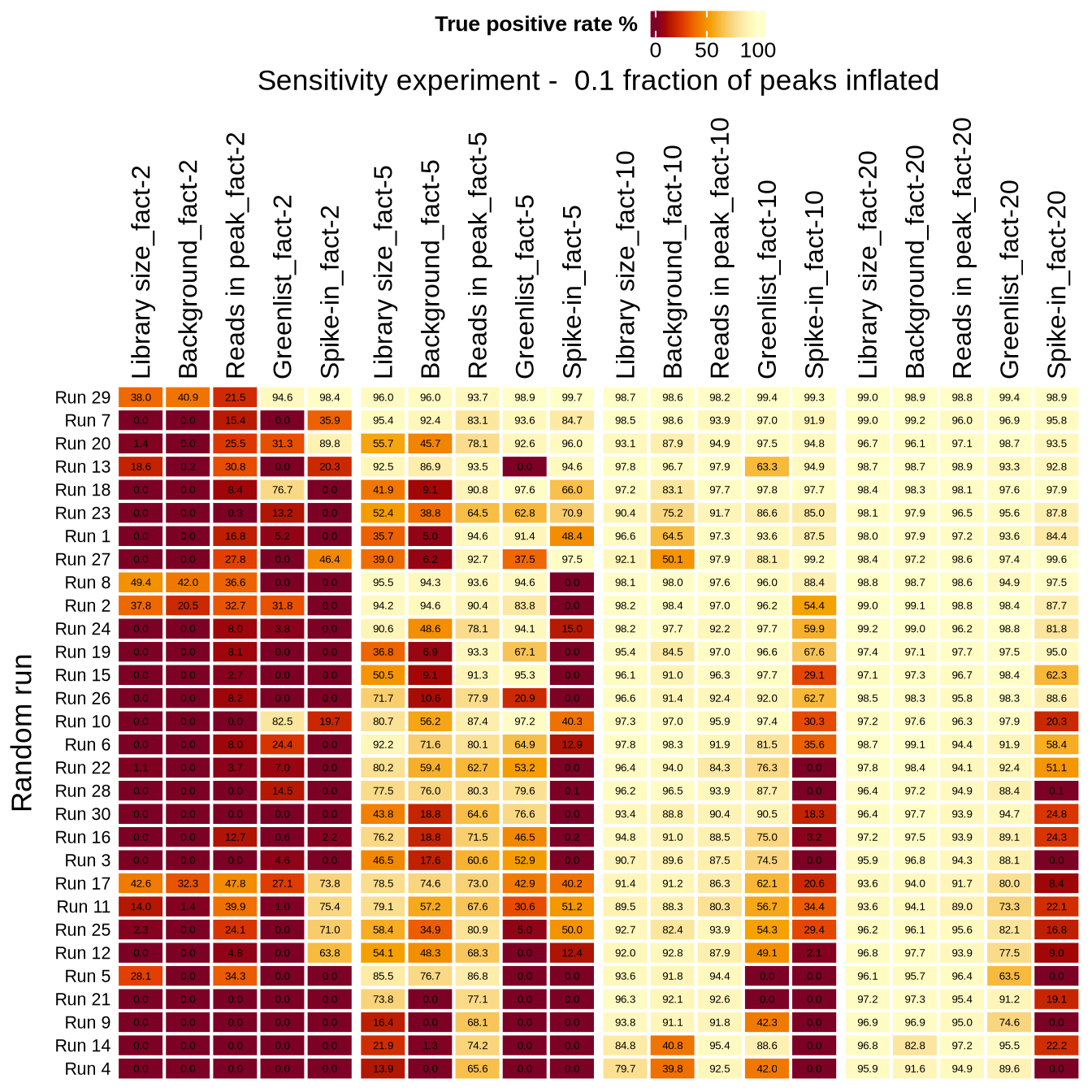

Sensitivity analysis of the global shift experiment by run when 10% of peaks were inflated. Inflation factors are given in the column names.

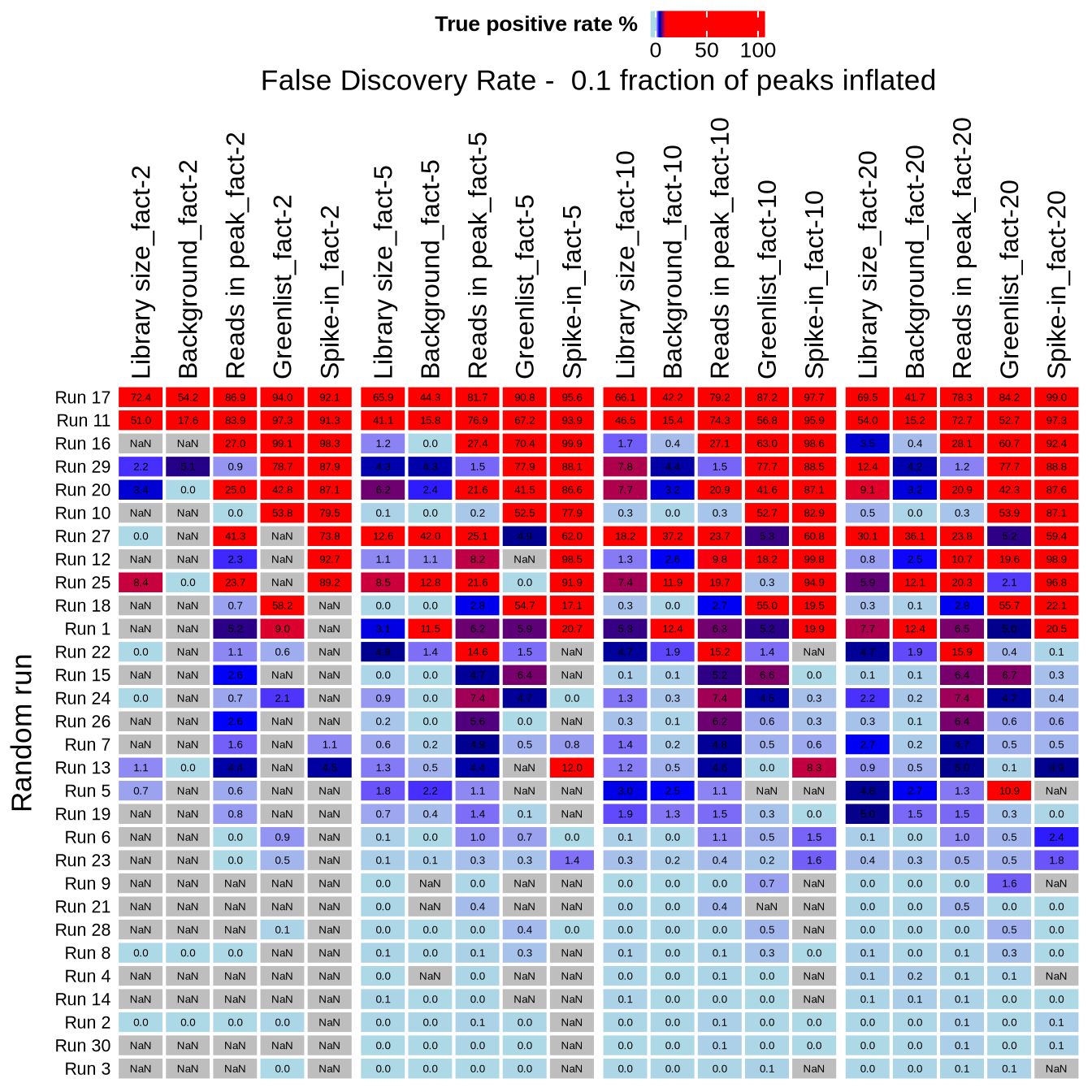

False discovery rate of the global shift experiment by run when 10% of peaks were inflated. Inflation factors are given in the column names.

### Supplementary Figure 2C: Global shift for 0.5 of peaks - sensitivity and FDR.

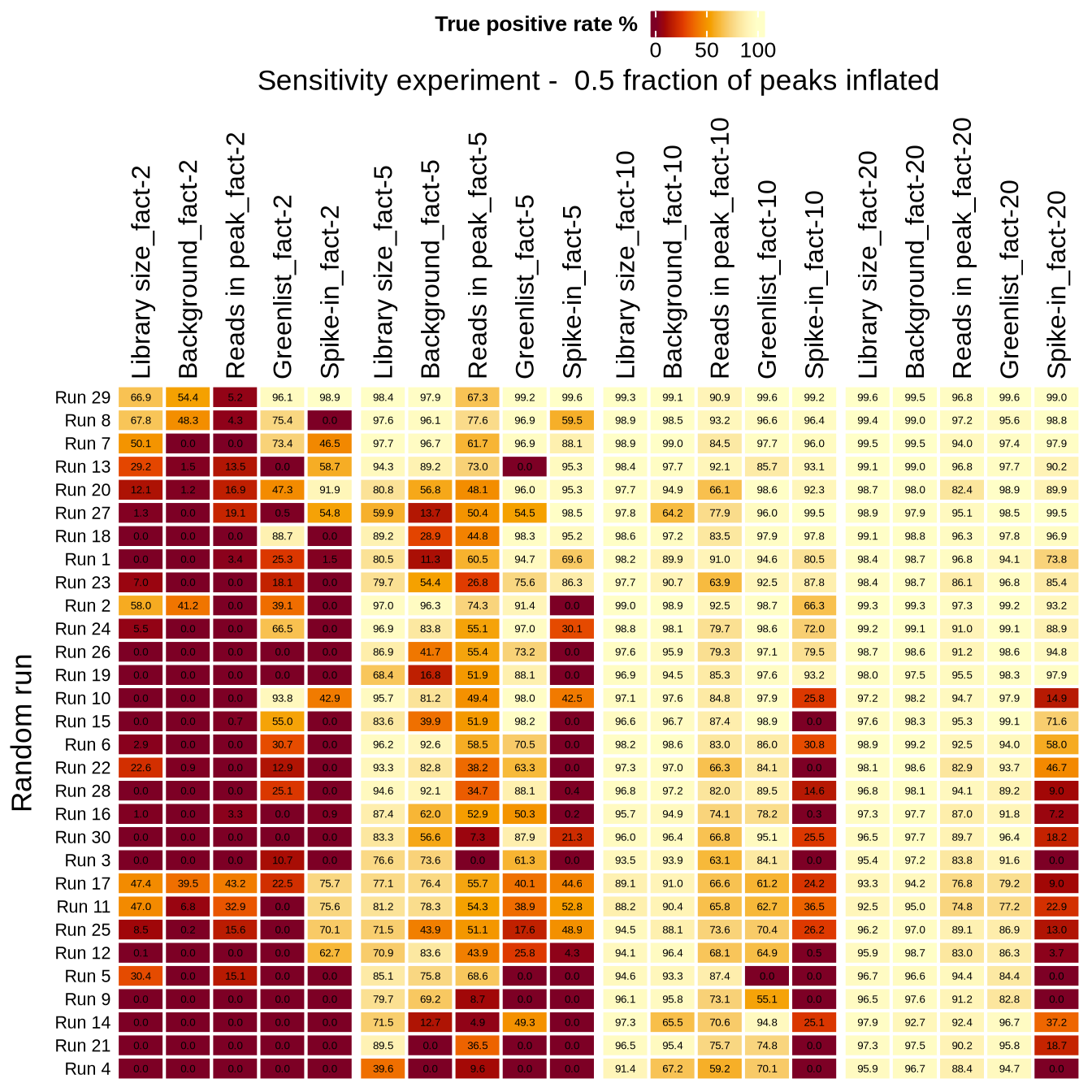

Sensitivity analysis of the global shift experiment by run when 50% of peaks were inflated. Inflation factors are given in the column names.

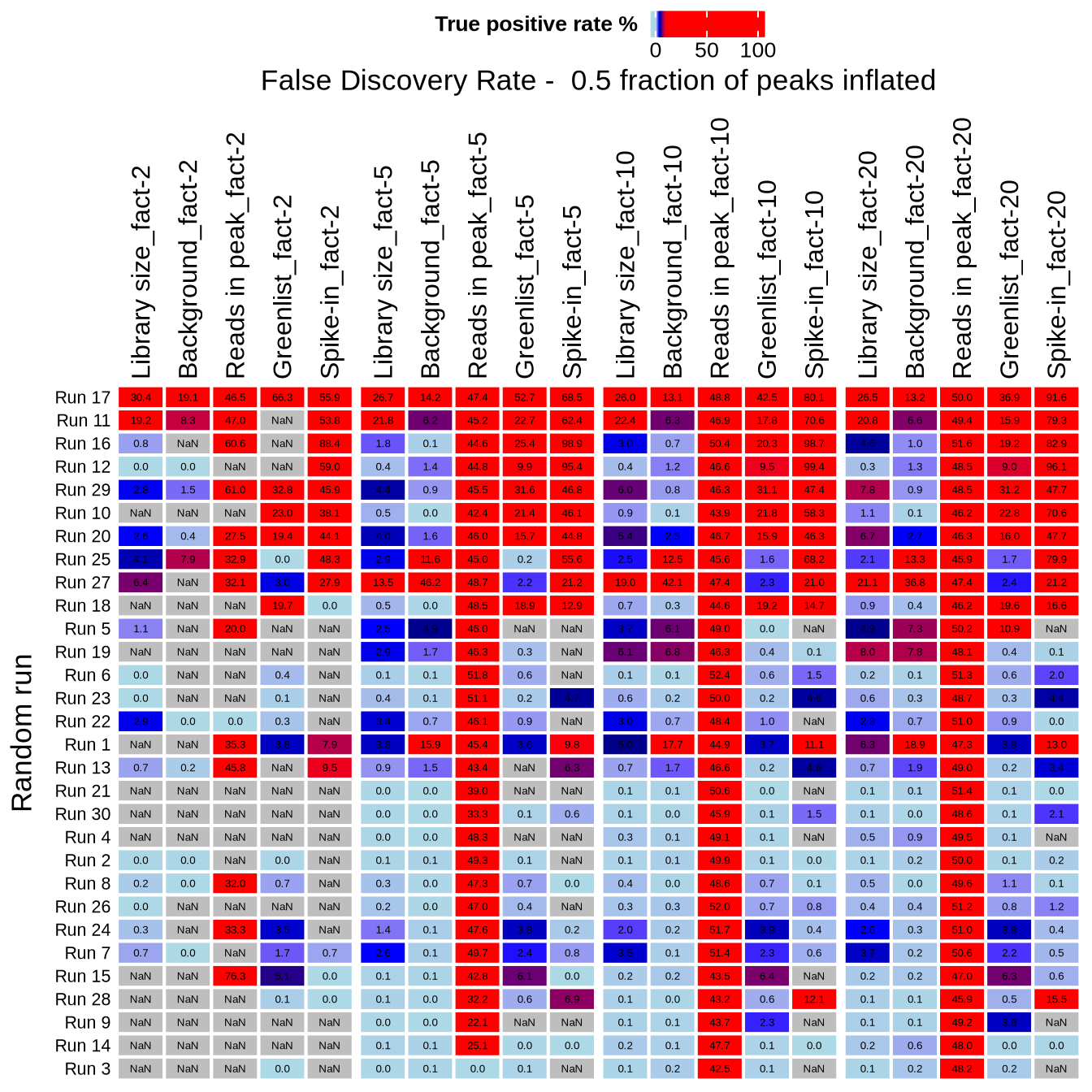

False discovery rate of the global shift experiment by run when 50% of peaks were inflated. Inflation factors are given in the column names.

### Supplementary Figure 2D: Global shift for 0.7 of peaks - sensitivity and FDR.

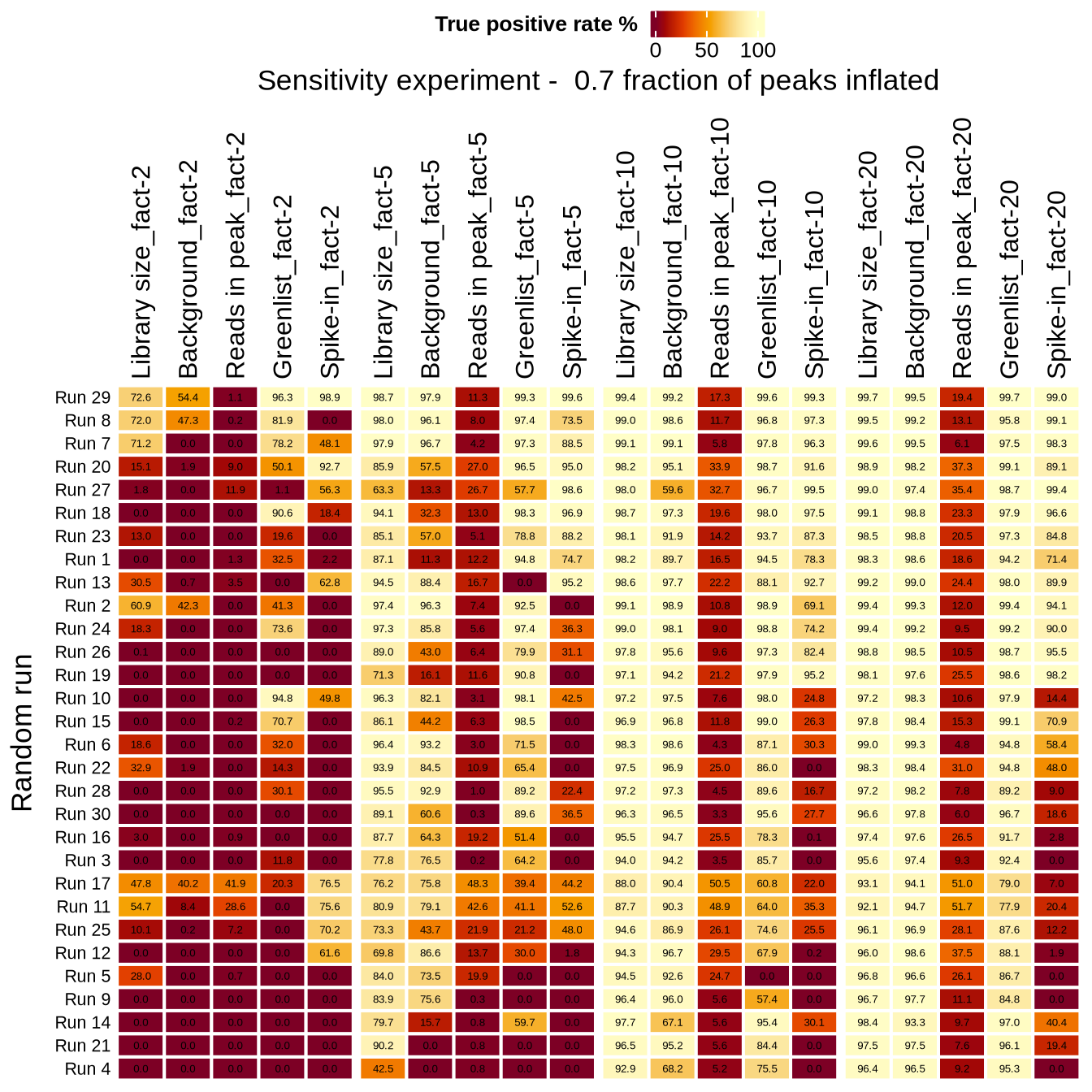

Sensitivity analysis of the global shift experiment by run when 70% of peaks were inflated. Inflation factors are given in the column names.

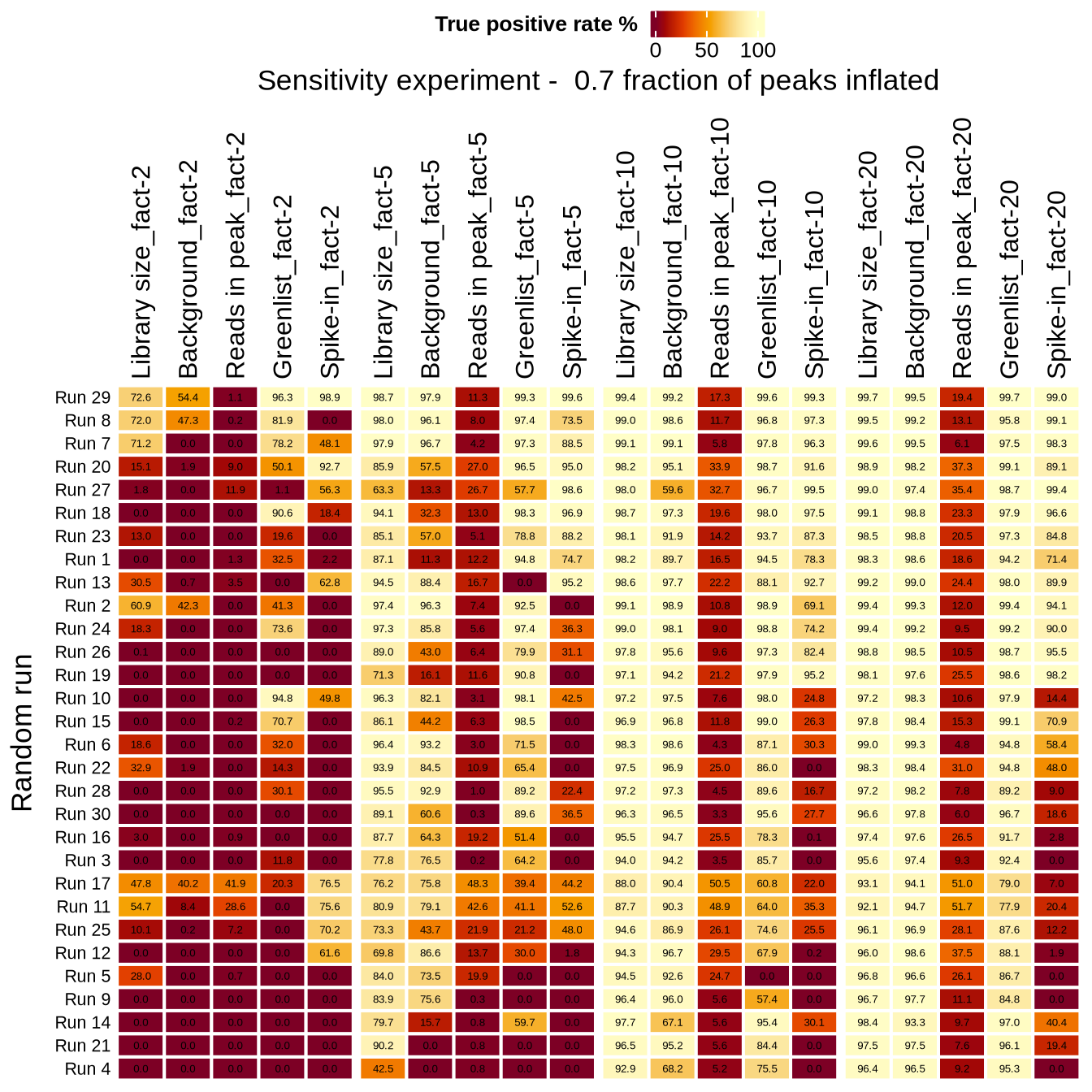

False discovery rate of the global shift experiment by run when 70% of peaks were inflated. Inflation factors are given in the column names.

### Supplementary Figure 3A: ER dataset - Insert size distribution

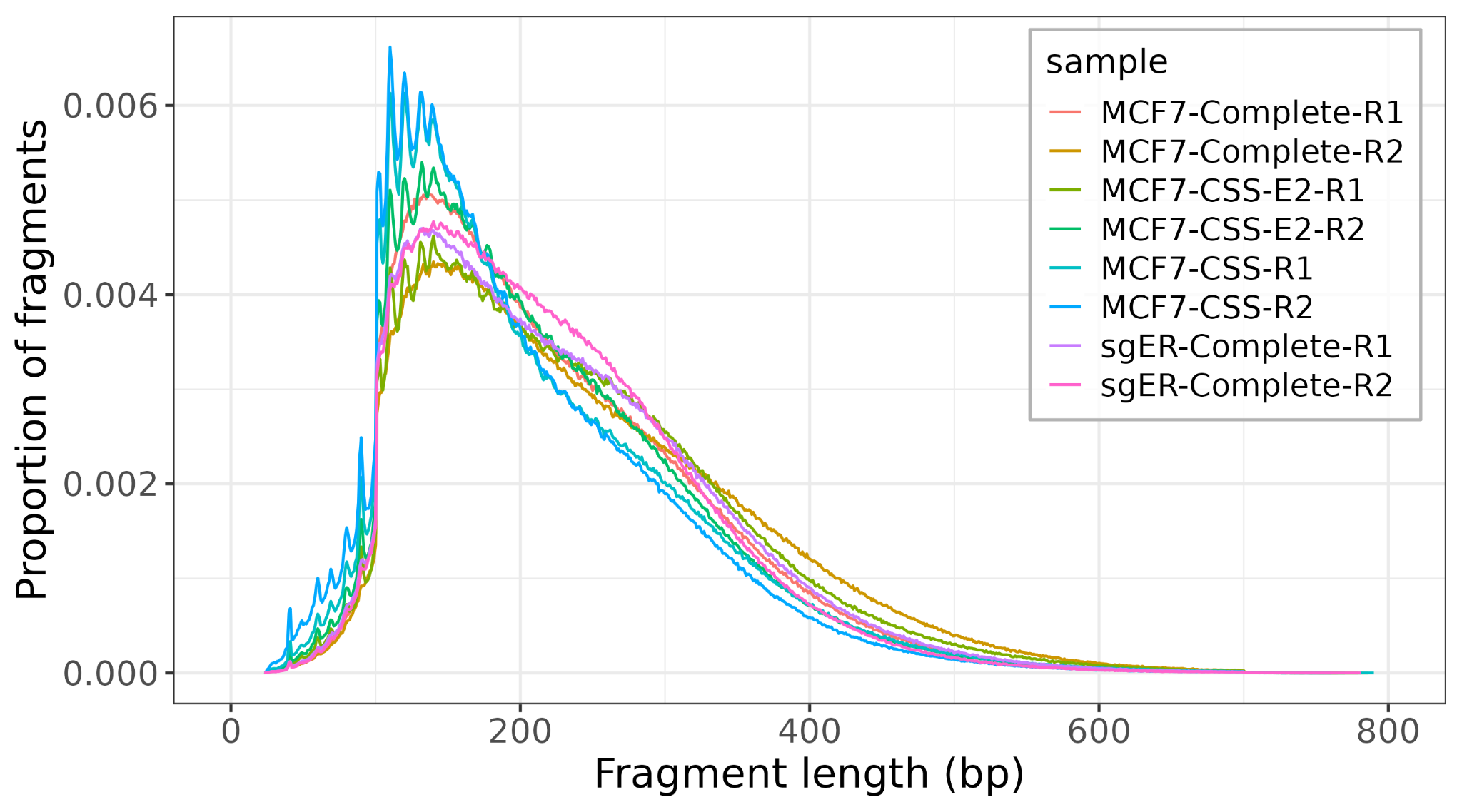

### Supplementary Figure 3B: ER dataset - raw signal read-enrichment heatmap

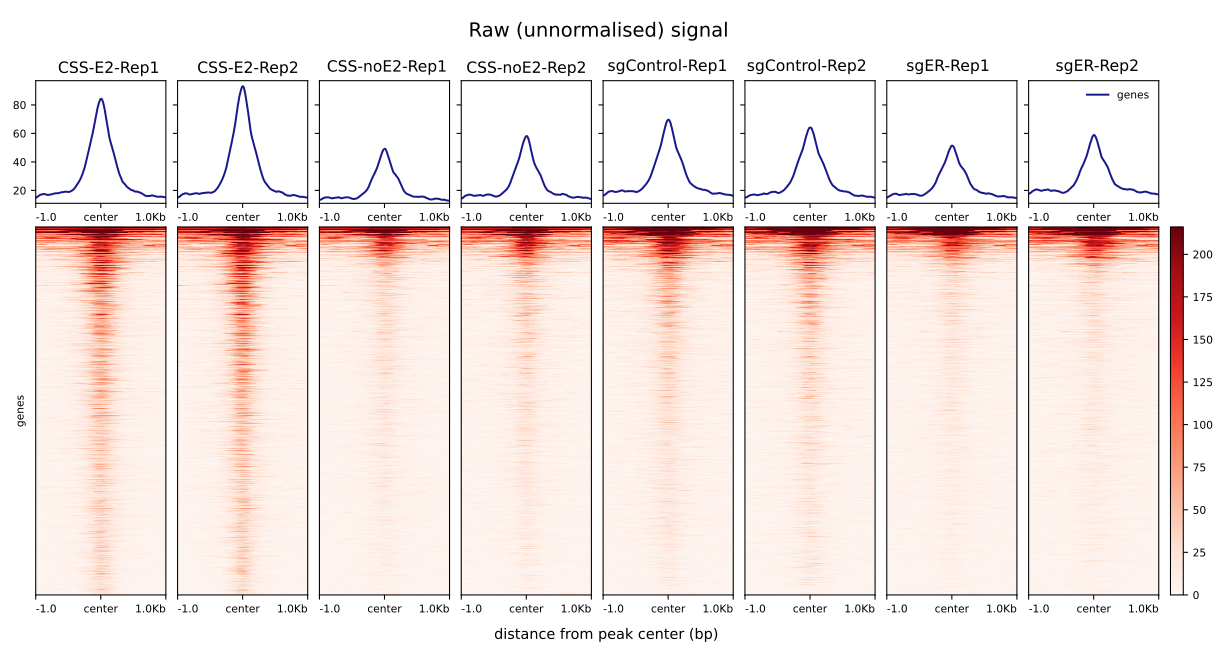

### Supplementary Table 3: ER dataset - aligned reads, spike-in scale factor and peak counts.

|  | BAM counts | | | | Duplication rate (%) | spike-in scale factor | | MACS2 peak counts with input |
| --- | --- | --- | --- | --- | --- | --- | --- | --- |
| Sample | hg38_dm6_sacCer3 | hg38 | dm6 | sacCer3 | hg38 | dm6 | sacCer3 |  |
| E2  Rep 1 | 27,699,128 | 23,162,887 | 1,963,124 | 1,428,886 | 31.54% | 0.509 | 0.699 | 1424 |
| E2  Rep 2 | 27,658,298 | 23,514,664 | 1,821,550 | 1,211,764 | 35.59% | 0.548 | 0.825 | 1834 |
| no E2 Rep 1 | 27,080,986 | 23,351,789 | 1,501,888 | 1,055,384 | 34.26% | 0.665 | 0.947 | 491 |
| no E2 Rep 2 | 32,062,746 | 27,510,077 | 1,790,908 | 1,315,286 | 42.10% | 0.558 | 0.76 | 710 |
| sg Control Rep 1 | 30,289,008 | 27,009,263 | 1,373,642 | 1,120,834 | 38.36% | 0.727 | 0.892 | 739 |
| sg Control Rep 2 | 29,017,464 | 24,820,688 | 1,881,358 | 1,592,030 | 42.79% | 0.531 | 0.628 | 439 |
| sgER Rep 1 | 27,021,674 | 24,129,233 | 1,107,518 | 1,057,066 | 32.13% | 0.902 | 0.946 | 204 |
| sgER Rep 2 | 30,259,592 | 27,286,431 | 1,051,564 | 1,095,556 | 35.00% | 0.95 | 0.912 | 252 |

Overview of samples; number of reads aligned to the hybrid genome, and to the individual species. Reads aligning to the hg38 blacklisted regions were removed. Duplicate rates (marked by Picard) of aligned hg38 BAM files. Scale factors determined by spike-in counts are given. The number of peaks called by MACS2 using the input sample in the “-control” parameter.

### Supplementary Figure 4: Western Blot validating ER knockdown

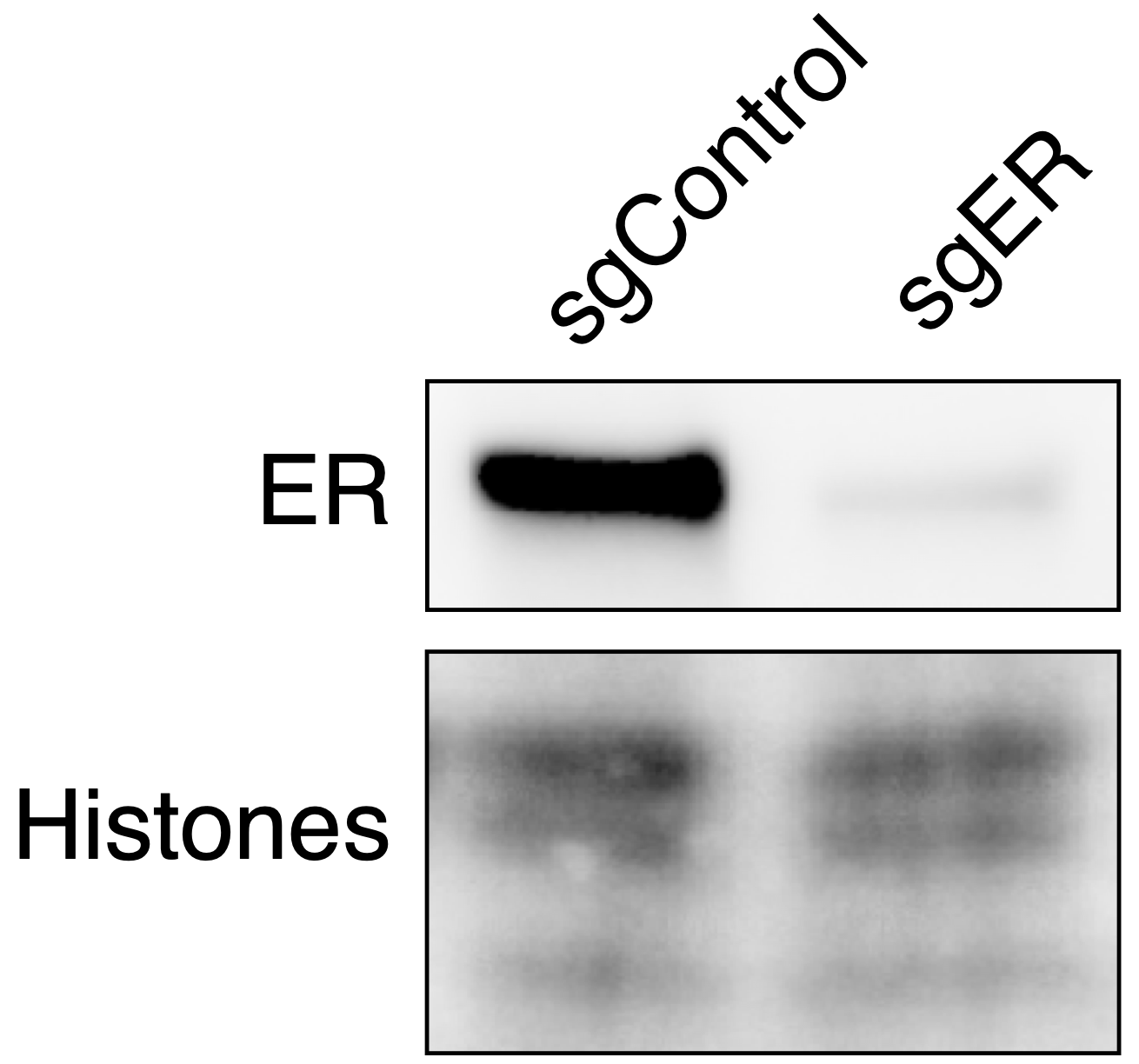

Western blot validating the ER knockdown in MCF7 cells used in ER CUT&RUN dataset.

### Supplementary Figure 5: PCA plot of ER datasets before normalisation

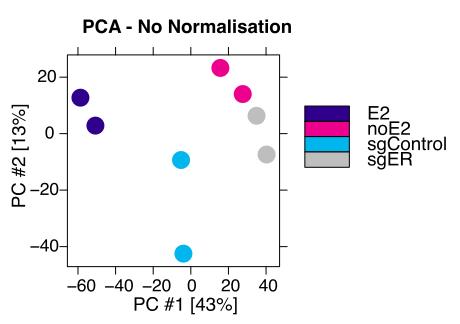

Created using: dbObj_counts_summit %>% dba.plotPCA(.,attributes = DBA_TREATMENT, score=DBA_SCORE_READS)

### Supplementary Figure 6: PCA plot of ER datasets with different normalisation methods

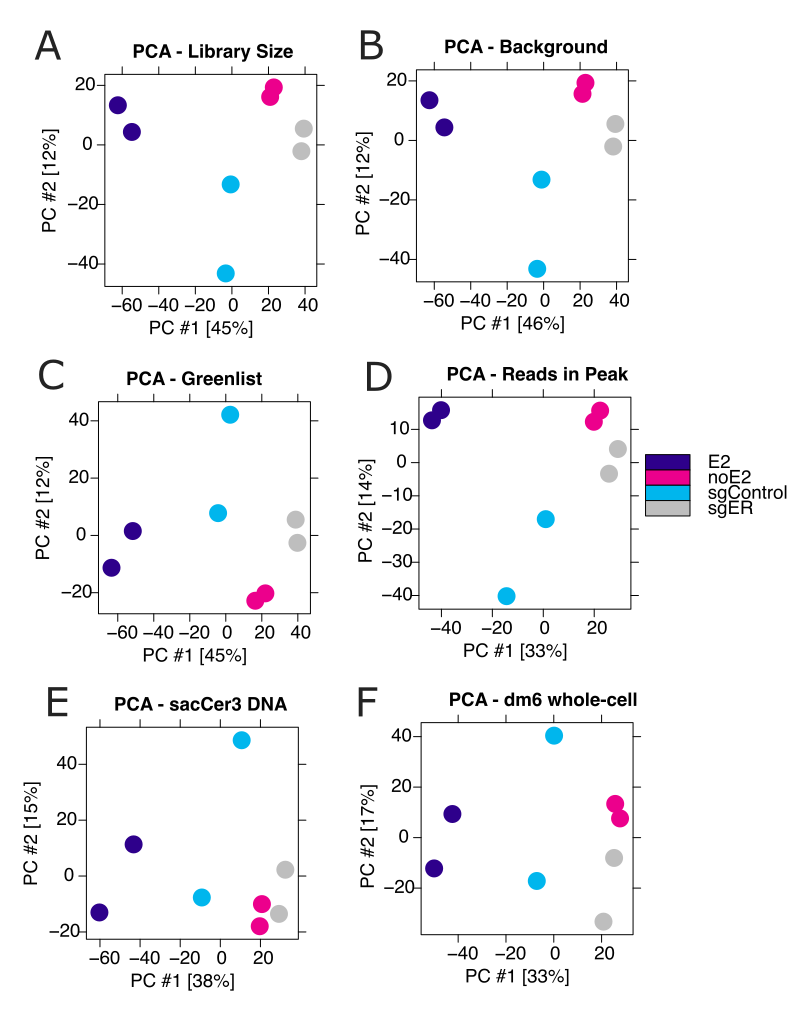

A - PCA of samples normalised by library size.

B - PCA of samples normalised by background.

C - PCA of samples normalised by greenlist.

D - PCA of samples normalised by reads in peak

E - PCA of samples normalised by sacCer DNA.

F - PCA of samples normalised by dm6 whole-cell.

### Supplementary Figure 7: Read-enrichment heatmaps of significant peaks unique to library size and background methods, not found using spike-in methods

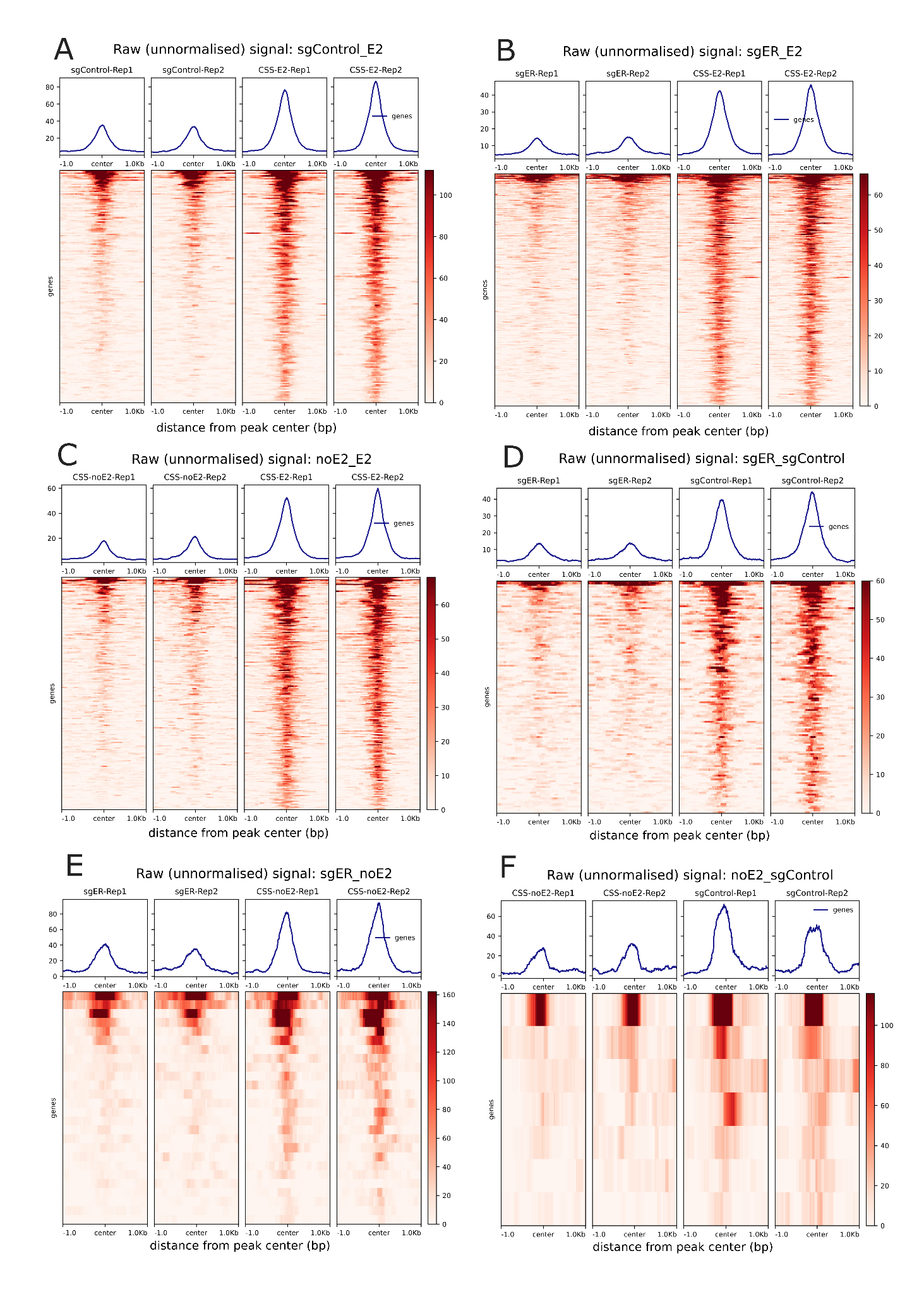

### Supplementary Figure 8: Cistrome of peaks unique to library size and background methods, not found in spike-in methods

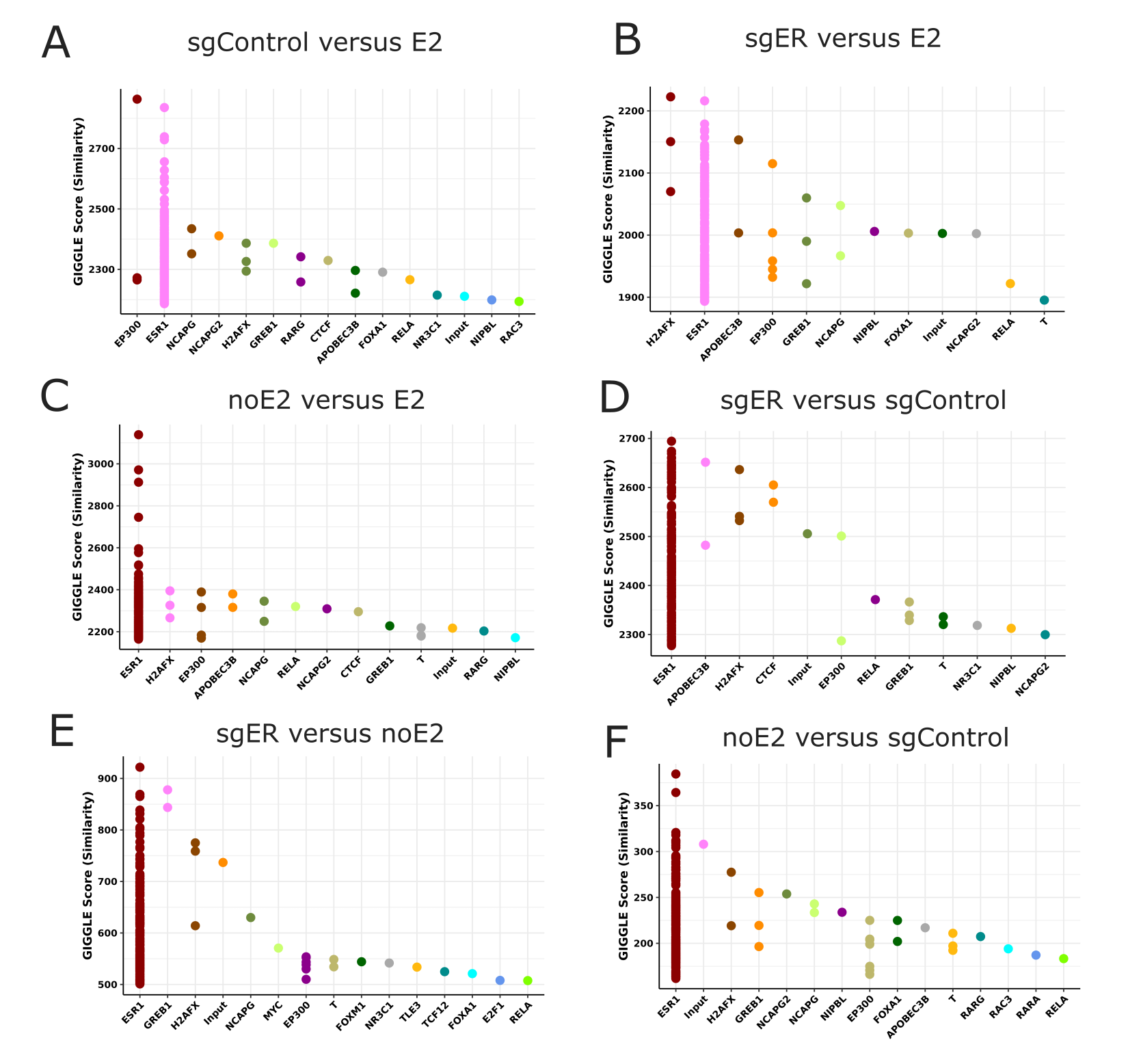

### Supplementary Table 4: BRG1 - aligned reads, peak numbers and fraction of reads in peak

|  | BAM counts | | | MACS 2 peak counts with input | fraction of reads in peaks |
| --- | --- | --- | --- | --- | --- |
| basename | hg38_sacCer3 | hg38 | sacCer3 |  |  |
| DMSO Rep 1 | 26,378,112 | 25,678,826 | 9,752 | 19,291 | 0.07 |
| DMSO Rep 2 | 25,894,752 | 25,218,508 | 13,046 | 23,001 | 0.1 |
| Abema Rep 1 | 19,881,888 | 19,285,951 | 5,226 | 21,769 | 0.07 |
| Abema Rep 2 | 27,185,982 | 26,336,707 | 9,564 | 22,168 | 0.07 |

Overview of samples; number of reads aligned to the hybrid genome, and to the individual species. Reads aligning to the hg38 blacklisted regions were removed. The fraction of reads in peak is given by DiffBind and refers to the reads overlapping the consensus peakset.

### Supplementary Figure 9: BRG1 dataset Quality Check

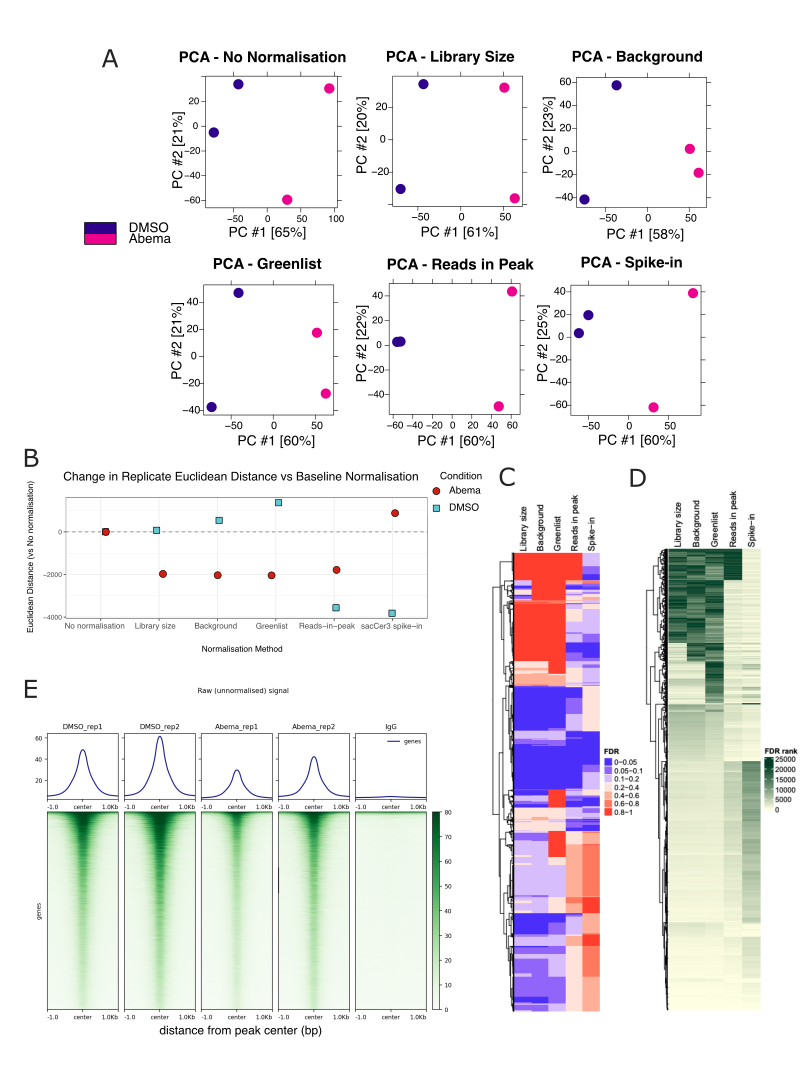

A. PCA plots by normalisation method, including before normalisation
B. Change in Euclidean distance between replicates compared to baseline (no normalisation) for each normalisation method
C. FDR value by normalisation method for significant differentially bound peaks that were not selected by all methods
D. FDR ranking by normalisation method for significant differentially bound peaks that were not selected by all methods
E. Raw unnormalised signal from BAM files

### Supplementary Figure 9F: BRG1 dataset insert size distribution

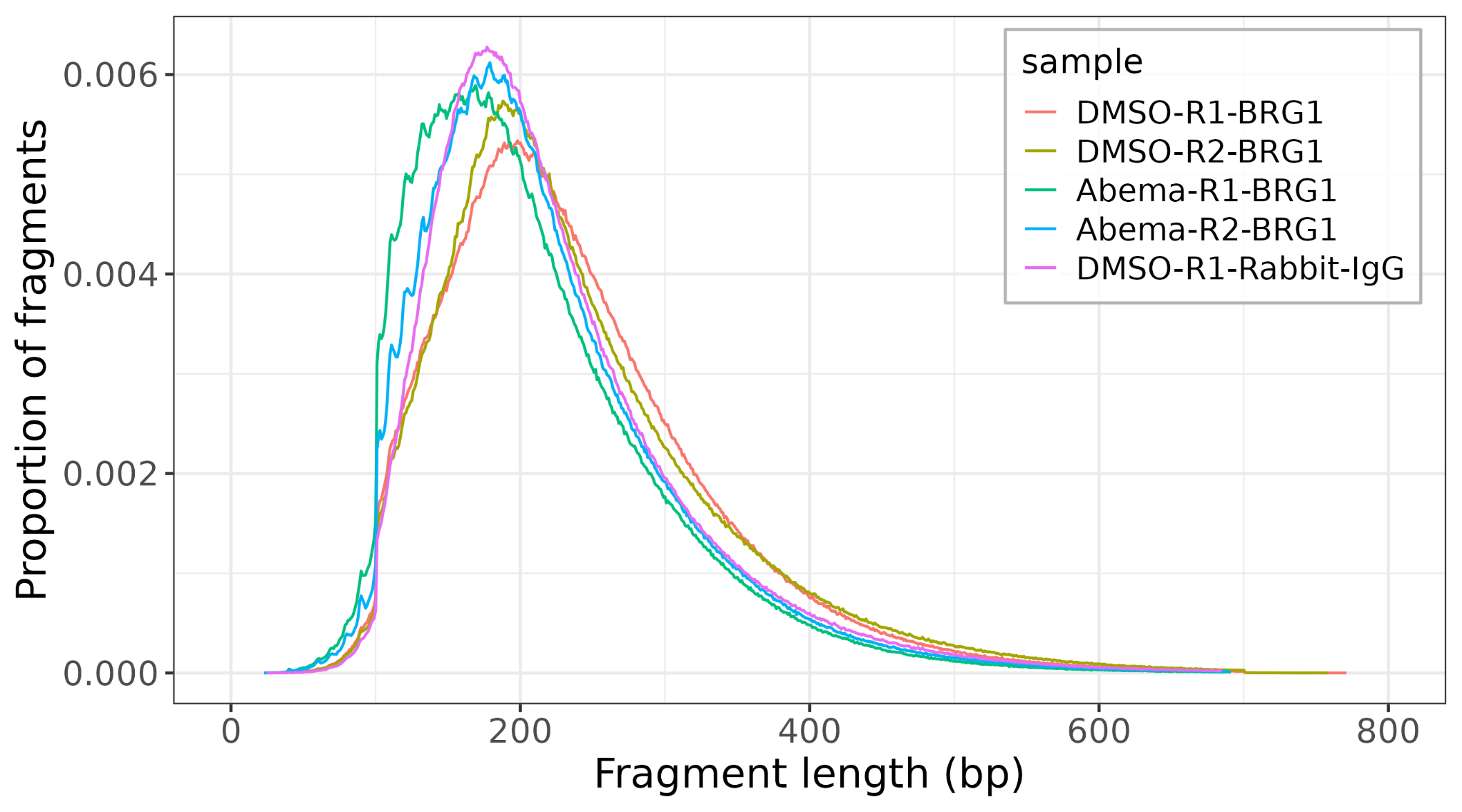

### Supplementary Figure 9G: BRG1 western blot

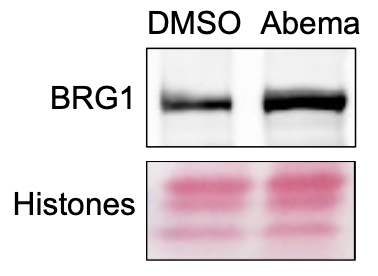

Samples were loaded by cell/nucleus number.
